# Mitochondrial folate metabolism inhibition drives differentiation through mTORC1 mediated purine sensing

**DOI:** 10.1101/2022.12.21.521404

**Authors:** Martha M. Zarou, Kevin M. Rattigan, Daniele Sarnello, Amy Dawson, Angela Ianniciello, Karen Dunn, Mhairi Copland, David Sumpton, Alexei Vazquez, G. Vignir Helgason

## Abstract

Supporting cell proliferation through nucleotide biosynthesis is an essential requirement for cancer cells. Hence, inhibition of folate-mediated one carbon (1C) metabolism, which is required for nucleotide synthesis, has been successfully exploited in anti-cancer therapy. Here, we reveal that mitochondrial folate metabolism is upregulated in patient-derived leukaemic stem cells (LSCs). We demonstrate that inhibition of mitochondrial 1C metabolism through impairment of *de novo* purine synthesis has a cytostatic effect on chronic myeloid leukaemia (CML) cells. Consequently, changes in purine nucleotide levels lead to activation of AMPK signalling and suppression of mTORC1 activity. Notably, suppression of mitochondrial 1C metabolism increases expression of erythroid differentiation markers. Moreover, we find that increased differentiation occurs independently of AMPK signalling and can be reversed through reconstitution of purine levels and reactivation of mTORC1. Of clinical relevance, we identify that combination of 1C metabolism inhibition with imatinib, a frontline treatment for CML patients, decreases the number of therapy-resistant CML LSCs in a patient-derived xenograft model. Our results highlight a novel role for folate metabolism and purine sensing in stem cell fate decisions and leukaemogenesis.

## Introduction

Rapidly dividing cells require building blocks to sustain growth and proliferation. Folate metabolism, through a series of complementary cytosolic and mitochondrial reactions, provides 1C units for biosynthetic processes including nucleotide synthesis and the methionine cycle (Fig. 1a)^1, 2^. New 1C units are donated from amino acids such as serine, the primary 1C unit donor, which gets converted into glycine via the enzyme serine hydroxymethyltransferase (SHMT)^1, 3, 4^. In parallel, activated 1C units are carried via folate molecules, allowing their assembly in different oxidative states^5^.

**Fig. 1:**
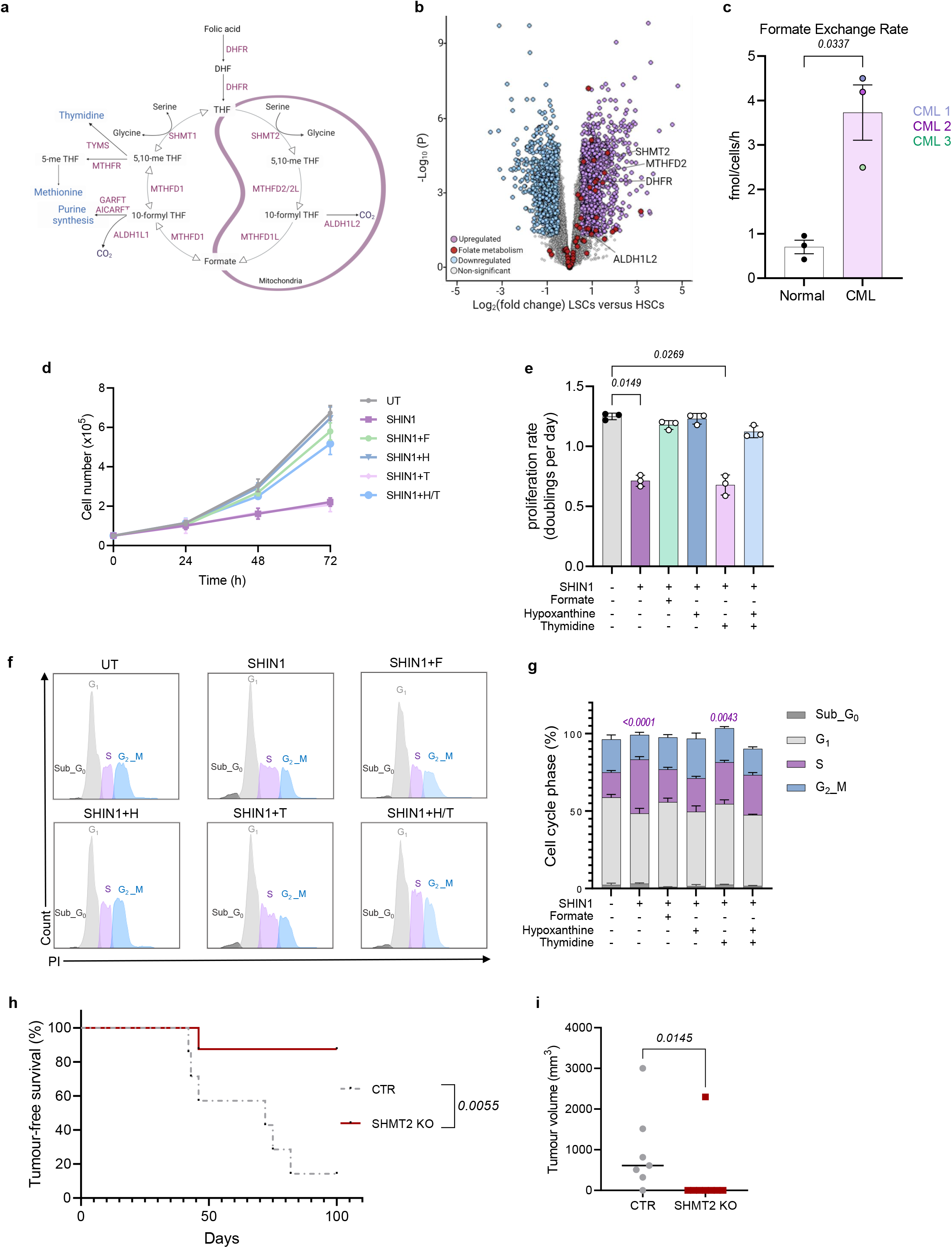
Folate metabolism is deregulated in LSCs and necessary for tumour formation *in vivo*. **a**, Schematic showing the compartmentalised folate metabolism pathway. DHF, dihydrofolate; THF; tetrahydrofolate; DHFR, dihydrofolate reductase; SHMT, serine hydroxymethyl transferase; 5,10-meTHF, 5,10, methenyl THF; MTHFD, methylenetetrahydrofolate dehydrogenesase; ALDH1L, 10-formyltetrahydrofolate dehydrogenase; GARFT, phosphoribosyl glycinamide formyl transferase; AICARFT, aminoimidazole-4-carboxamide ribonucleotide formyl transferase, MTHFR, methylenetetrahydrofolate reductase; TYMS, thymidylate synthase; 5-meTHF, 5-methyl THF. **b**, Volcano plot of differentially expressed genes in CML LSCs compared to normal HSCs (CD34^+^CD38^-^) (E-MTAB-2581). Upregulated genes with a LSCs versus HSCs log2 (fold change) of at least 0.5 are shown in light purple, while downregulated gene with a LSCs versus HSCs log2 (fold change) of at least −0.5 are shown in blue. Folate metabolism associated genes are shown in red. **c**, Formate release rate from normal and CML CD34^+^ conditioned media following 48 h culture (n=3 patient samples). **d,e,** Growth (**d**) and proliferation rate (**e**) of untreated (UT) K562 or cultured with 2.5 μM SHIN1 with or without the addition of 1 mM formate (F), 100 μM hypoxanthine (H), 16 μM thymidine (T) for 72 h (n=3 independent cultures). **f,**Representative flow cytometry histograms obtained from cell cycle analysis of K562 cells treated as in (**d,e**) using propidium iodide (PI) staining. **g**, Percentage of cell cycle phases as depicted in (**f**) (n=3 independent cultures). **h,i,** Tumour-free survival (**h**) and tumour volume (**i**) at endpoint of NRGW^41^ mice transplanted with KCL22 control (CTR) (n=7 mice) and SHMT2 KO cells (n=8 mice). Data are shown as the mean ± s.e.m. P-values were calculated using unpaired two-tailed t-test with Welch’s correction (**c**), a repeated measure one-way ANOVA with Dunnett’s multiple comparisons test (**e**), two-way ANOVA with Tukey’s multiple comparison’s test (**f**) relative to untreated K562 cells, log-rank (Mantel-Cox) test for survival analysis (**g**) or unpaired Mann-Whitney test (**h**).

In agreement with its key role for nucleotide synthesis, mitochondrial SHMT2 and the immediately downstream 5,10-methylene-tetrahydrofolate dehydrogenase (MTHFD2), are consistently overexpressed in cancer^6, 7^. Furthermore, serine catabolism to formate is increased in intestinal adenomas and mammary carcinomas^8^. Reinforcing of the requirement for folate metabolism in cancer, antifolates have proven highly effective as chemotherapeutic agents. Methotrexate, the most recognised antifolate, has been used in the treatment of a variety of neoplasms and inflammatory diseases^9, 10^. Importantly, antifolates have found a wide application in haematological malignancies^11, 12^. However, the role of folate metabolism in LSCs remains undescribed.

CML is a stem cell-driven myeloproliferative disorder that arises in a single haematopoietic stem cell (HSC) with the introduction of the BCR-ABL oncoprotein^13^. BCR-ABL, through activation of several molecular pathways including RAS, MYC and PI3K/AKT/mTOR, is the main driver of leukaemia initiation and progression^14–16^. As exemplified in CML, increasing evidence suggests that stem cells, but not fully differentiated cells, are a major driver of neoplasms and therapy resistance^17–19^. Oncogenic stem cells evade standard therapy due to their quiescent/slow-cycling status^20^. To maintain tissue homeostasis, stem cells, including HSCs, are characterised by their capacity to self-renew and their ability to differentiate^21, 22^. Thus, identification of vulnerabilities that when targeted, can stimulate differentiation of stem/progenitor cell population can have potent anticancer effects^23–25^.

Strikingly, slow-growing CML LSCs have been found to resist treatment with tyrosine kinase inhibitors (TKIs), even when BCR-ABL1 kinase activity is fully inhibited^26–28^. It has been established that BCR-ABL1 rewires metabolic programmes such as oxidative phosphorylation (OXPHOS) and the tricarboxylic acid cycle^29, 30^. Intriguingly, targeting OXPHOS in combination with standard therapy was found to eradicate persistent LSCs^29^. Furthermore, it was recently demonstrated that exogenous serine controls the cell fate of epidermal stem cells during tumour initiation^31^. These studies highlight how metabolic rewiring of oncogenic stem cells might affect tumour initiation and persistence. In the present study, we describe deregulation of mitochondrial folate metabolism in leukaemic stem and progenitor cells. We demonstrate that inhibition of mitochondrial folate metabolism through impairment of *de novo* purine synthesis drives differentiation of CML cells. Mechanistically, we show that the differentiation following nucleotide depletion depends on suppression of mTORC1 activity through decreased purine sensing. Moreover, we uncover that inhibition of 1C metabolism sensitises human LSCs to standard CML therapy. Collectively, our results provide novel insight into the role of folate metabolism in critical cell fate decisions.

## Results

### Mitochondrial folate metabolism is deregulated in LSCs and necessary for tumour formation *in vivo*

To assess whether folate metabolism is deregulated in primitive CML cells, we carried out differential gene expression analysis of a transcriptomic dataset generated from CML LSCs (CD34^+^CD38^-^) and normal HSCs (CD34^+^CD38^-^)^32^. Gene Set Enrichment Analysis (GSEA) revealed that folate metabolism is upregulated in LSCs (Extended Data Fig. 1a,b). Specifically, there was an upregulation of gene transcripts encoding enzymes involved in mitochondrial 1C reactions such as *ALDH1L2, SHMT2* and *MTHFD2* (Fig. 1b and Extended Data Fig. 1c). The upregulation of mitochondria-related genes supports the observation that most cancer cells produce 1C units exclusively in the mitochondria^5^. To confirm that folate metabolism is deregulated in primitive CML cells, we functionally determined the activity of folate metabolism in patient-derived CML CD34^+^ cells and normal counterparts. To this end, using gas chromatography-mass spectrometry (GS-MS) we assessed formate overflow as a proxy for the rate of serine catabolism. CML CD34^+^ cells had an increased formate exchange rate and extracellular formate concentration compared to normal CD34^+^ cells, confirming a higher rate of folate metabolism in CML cells (Fig. 1c and Extended Data Fig. 1d).

We next analysed the consequences of 1C metabolism inhibition in CML K562 cells. We used the pharmacological SHMT1/2 inhibitor SHIN1 (which inhibits cytosolic and mitochondrial arms of the pathway)^33^, in parallel with a genetic knockout (KO) of mitochondrial SHMT2 (Extended Data Fig. 1e). Due to the reversibility of the cytosolic compartment of folate metabolism, mitochondrial 1C metabolism is believed to be dispensable^5, 34^. As such, single deletions of mitochondrial enzymes do not consistently block cancer cell growth in nutrient rich conditions^5, 34, 35^. Strikingly, we observed a proliferation halt following either pharmacological (cytosolic/mitochondrial) or genetic (mitochondrial) inhibition of 1C metabolism (Fig. 1d,e and Extended Data Fig. 1f,g). Furthermore, SHIN1 treatment or SHMT2 KO resulted in accumulation of K562 cells in the S phase of the cell cycle (Fig. 1f,g and Extended Data Fig. 1h). Formate supplementation was sufficient to restore growth, indicating that 1C units are the limiting factor for proliferation and cell cycle progression of CML cells (Fig. 1d-g and Extended Data Fig. 1f-h). To assess whether purine or thymidylate depletion caused the proliferation arrest following inhibition of 1C metabolism, we supplemented cells with the purine salvage precursor hypoxanthine, thymidine or a combination of both. While hypoxanthine rescued proliferation, thymidine did not (Fig. 1d-g and Extended Data Fig. 1f-h). These data are consistent with the fact that purine biosynthesis requires a higher flux of 1C units compared to thymidylate synthesis^36^. It is likely that in the short-term, residual 1C units can sustain thymidylate synthesis. Our findings support that nucleotide impairment following pharmacological or genetic inhibition of folate metabolism has a cytostatic effect on CML cells.

CML KCL22 cells form extramedullary tumours when transplanted in immunocompromised mice. To test whether loss of the folate cycle impairs tumour formation *in vivo*, KCL22 SHMT2 KO cells were transplanted via tail vain injection into non-irradiated immunocompromised mice. Loss of SHMT2 significantly prolonged tumour-free survival of xenografted mice, when compared with mice receiving SHMT2 competent control cells (Fig. 1h). This correlated with a significant decrease in tumour burden in mice transplanted with KCL22 SHMT2 KO cells (Fig. 1i). Together, these results suggest that mitochondrial 1C metabolism sustains proliferation of CML cells both *in vitro* and *in vivo*, and cytosolic 1C metabolism does not compensate for the loss of SHMT2 in CML cells.

### Inhibition of mitochondrial serine catabolism disrupts *de novo* purine synthesis and glycolysis in CML cells

Following the observation that mitochondrial 1C metabolism inhibition results in proliferation arrest due to impairment of purine biosynthesis, we sought to investigate the downstream metabolic effects of folate cycle suppression. As expected, metabolic analysis revealed that SHIN1 treatment or SHMT2 deficiency led to reduction of the glycine pool (Fig. 2a and Extended Data Fig. 2a). Moreover, formate supplementation resulted in a further decrease of glycine levels, as it restores the 1C unit pool and drives *de novo* purine synthesis bypassing SHMT1/2 activity. Notably, we observed a dramatic increase in the purine biosynthetic intermediate 5′-phosphoribosyl-5-aminoimidazole-4-carboxamide (AICAR), and a corresponding reduction in purine nucleotides (Fig. 2c and Extended Data Fig. 2c). Changes in purine nucleotide pools were reversed by formate. Surprisingly, we observed a reduction in the ribose-5-phosphate (R5P) pool, which is upstream of a step requiring glycine. R5P, apart from being an intermediate of purine/pyrimidine synthesis, is a product of the non-oxidative phase of the pentose phosphate pathway (Fig. 2b and Extended Data Fig. 2b). Prompted by this reduction, we decided to investigate the effect of 1C metabolism inhibition on glycolysis related metabolites. Pharmacological or genetic inhibition of 1C metabolism resulted in increased intracellular glucose levels and decreased pyruvate and lactate levels (Fig. 2d and Extended Data Fig. 2d). Subsequently, quantitative analysis of the media from SHIN1-treated or SHMT2 KO cells confirmed a significant reduction in glucose uptake and lactate secretion, when compared with media from 1C metabolism competent cells (Fig. 2e and Extended data Fig. 2e). To confirm the decrease in glycolytic flux, we measured the extracellular acidification rate (ECAR) following suppression of 1C metabolism. SHIN1 treatment or SHMT2 ablation reduced both glycolysis and glycolytic capacity (Fig. 2f,g and Extended Data Fig. 2f,g). Of note, formate supplementation reversed the decrease in glycolytic flux, suggesting that impaired glycolysis was linked to reduced nucleotide levels. This is in agreement with our previous work where we applied theoretical modelling to indicate that 1C metabolism promotes glycolysis^37^.

**Fig. 2:**
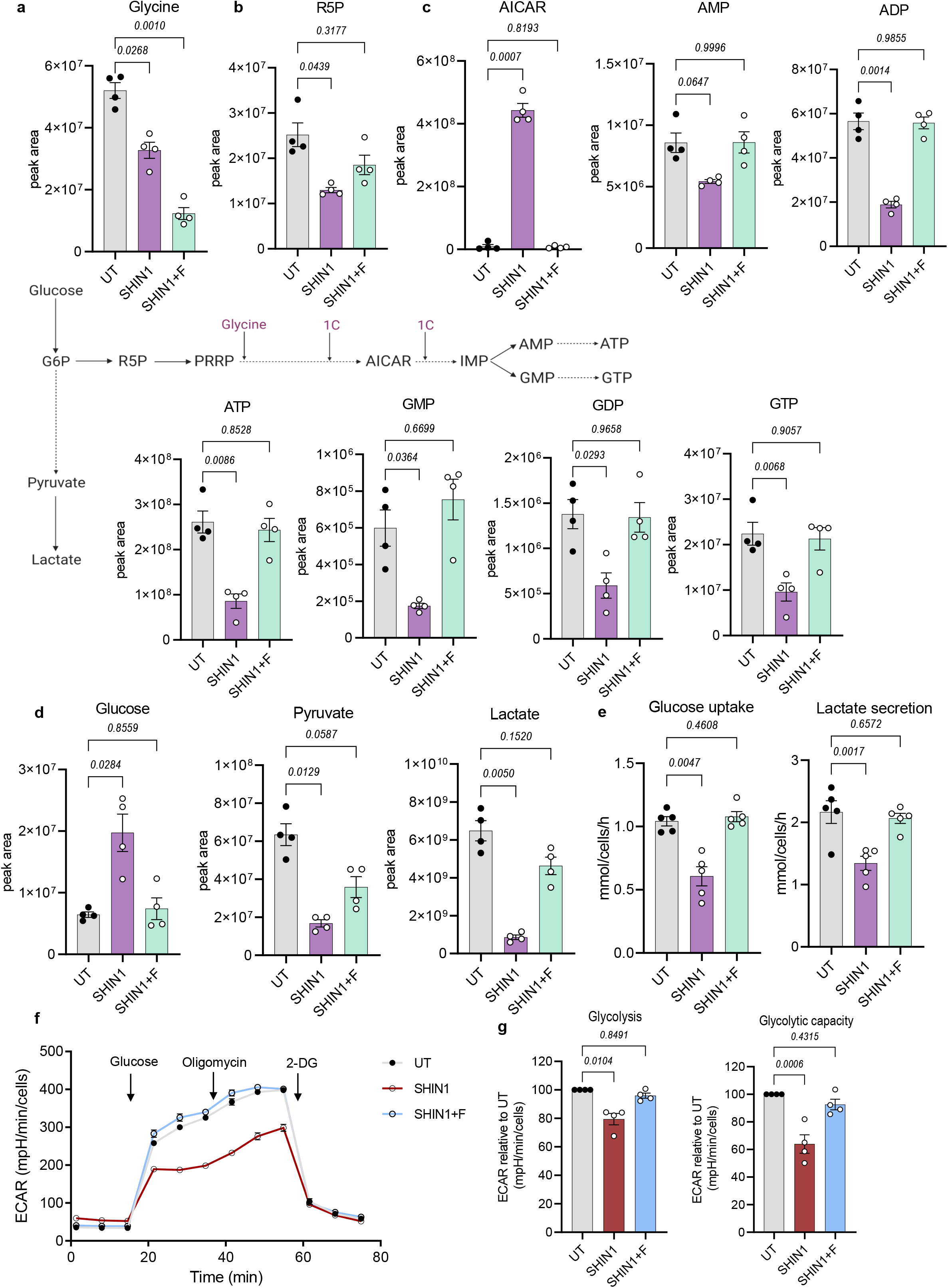
Inhibition of mitochondrial serine catabolism disrupts *de novo* purine synthesis and glycolysis in CML cells. **a-d,** LC-MS measurement of intracellular glycine (**a**), ribose-5-phosphate (R5P) (**b**), purine biosynthetic intermediate and nucleotides (**c**) and glycolysis related metabolites (**d**) in K562 cells treated with 2.5 μM SHIN1 with or without the addition of 1 mM formate for 24 h (n=4 independent cultures). **e**, Glucose uptake and lactate secretion in K562 cells in response to treatment as explained in (**a-d**) (n=5 independent cultures). **f,g,** Representative extracellular acidification rate (ECAR) profile (**f**), relative glycolysis and glycolytic capacity (**g**) as measured by the Seahorse XF analyser of K562 cells treated as in (**a-e**) (n=4 independent cultures). Data are shown as the mean ± s.e.m. P-values are derived from a repeated measure one-way ANOVA with Dunnett’s multiple comparisons test (**a-e**) or ordinary one-way ANOVA with Dunnett’s multiple comparisons test (**g**).

### Suppression of mitochondrial folate metabolism leads to differentiation of CML cells in an AMPK-independent manner

Since AICAR and purine nucleotides can directly or indirectly signal to AMP-activated protein kinase (AMPK)^38^, we next sought to investigate the effect of mitochondrial folate metabolism inhibition on AMPK activity. Indeed, SHIN1 treatment and SHMT2 KO induced AMPK activity, measured by increased phosphorylation of AMPK on threonine-172, and increased phosphorylation of the two AMPK effectors, acetyl-coA carboxylase (ACC) and Unc-51 like autophagy activating kinase 1 (ULK1) (Fig. 3a and Extended Data Fig. 3a). Increased ULK1-dependent phosphorylation of the autophagy related 13 (ATG13) protein suggests activation of the autophagy initiation complex (Fig. 3a and Extended Data Fig. 3a). Concurrently, pharmacological or genetic inhibition of folate metabolism reduced phosphorylation of the ribosomal protein S6 (RPS6), a target of the mechanistic target of rapamycin complex 1 (mTORC1) (Fig. 3a and Extended Data Fig. 3a). Treatment with AICAR resulted in a similar pattern of AMPK signalling activation and mTORC1 suppression, while combination of AICAR with folate metabolism inhibition did not have any additive effect (Fig. 3a and Extended Data Fig. 3a). Additionally, formate supplementation reversed the effect on AMPK and mTORC1 (Fig. 3a and Extended Data Fig. 3a).

**Fig. 3:**
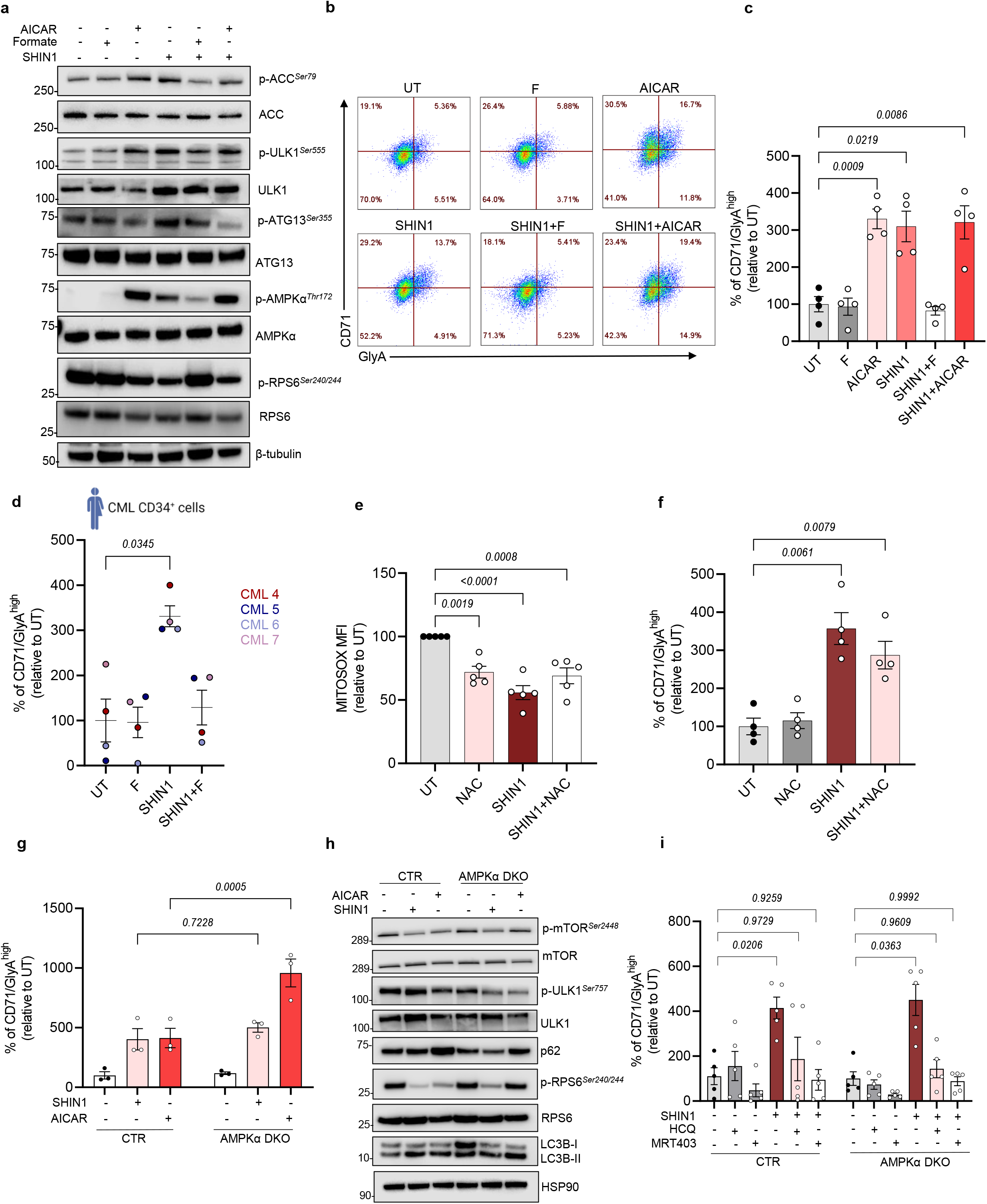
Suppression of mitochondrial folate metabolism leads to ROS and AMPK-independent differentiation of CML cells. **a**, Immunoblot analysis to assess AMPK and mTORC1 signalling in K562 cells following 24 h incubation with 1 mM AICAR, 2.5 μM SHIN1 in the presence or absence of 1 mM formate and combination of SHIN1 with AICAR (representative of three independent experiments). **b**, Representative flow cytometry plots of CD71 and GlyA expression in K562 cells subjected to treatment as in (**a**). **c**, Percentage of CD71/GlyA^high^ cells relative to untreated as depicted in (**b**) (n=4 independent cultures). **d**, Percentage of CD71/GlyA^high^ CML CD34^+^ cells relative to untreated after 72 h incubation with 2.5 μM SHIN1 with or without the supplementation of 1 mM formate. Cells were incubated in the presence of 3 U/ml of erythropoietin (n=4 patient samples). **e**, Quantification of relative mean fluorescent intensity (MFI) of mitochondrial ROS in K562 cells treated with 2.5 μM SHIN1 with or without the addition of 300 μM NAC for 72 h (n=4 independent cultures). **f**, Percentage of CD71/GlyA^high^ K562 cells relative to untreated after 72 h incubation with 2.5 μM SHIN1 in the presence or absence of 300 μM NAC (n=4 independent cultures). **g**, Relative percentage of CD71/GlyA^high^ K562 control and AMPKα DKO cells in the presence of 1 mM AICAR or 2.5 μM SHIN1 for 72 h (n=4 independent cultures). **h**, Western blot analysis to assess mTORC1 activity and autophagy induction in K562 control and AMPKα DKO cells following 24 h culture with the addition of 1 mM AICAR or 2.5 μM SHIN1 (representative of three independent experiments). **i**, Relative percentage of CD71/GlyA^high^ K562 control and AMPKα DKO cells treated with 2.5 μM SHIN1 with or without the addition of 5 μM hydroxychloroquine (HCQ) for 72 h (n=5 independent cultures). Data are shown as the mean ± s.e.m. P-values were calculated with a repeated measure one-way ANOVA with Dunnett’s multiple comparisons test (**c,f,i**), Kruskal-Wallis test with Dunn’s multiple comparisons test (**d**), ordinary one-way ANOVA with Dunnett’s multiple comparisons test (**e**) or two-way ANOVA with Sidak’s multiple comparisons test (**g**).

In addition to being an activator of AMPK, AICAR has been shown to promote differentiation of acute myeloid leukaemia (AML) cells^39, 40^. Thus, we next assessed whether inhibition of mitochondrial folate metabolism induces differentiation of CML K562 cells. These cells have been shown to be a suitable model to study differentiation, as they undergo erythroid maturation when treated with differentiation-inducing compounds^41^. Macroscopic examination revealed that cell pellets appeared with a strong red colour following inhibition of mitochondrial serine catabolism, which suggests a higher haemoglobin production and, ultimately, an increase in erythroid maturation (Extended Data 3b). To confirm initiation of erythroid maturation, we stained K562 cells for erythropoiesis markers: Glycophorin A (GlyA), a sialoglycoprotein found in the membrane of erythrocytes, and transferrin receptor 1 (CD71), a transmembrane glycoprotein necessary for iron import inside the cells^42^. SHIN1 treatment resulted in enhanced levels of GlyA and CD71 (Fig. 3b,c). Similarly, expression of GlyA and CD71 increased following loss of SHMT2 (Extended Data Fig. 3c). Of note, AICAR treatment promoted expression of GlyA and CD71 to the same extent as SHIN1, while we did not observe any additive effect when combining AICAR with SHIN1 (Fig. 3b,c). Restoration of the 1C unit pool through formate supplementation was sufficient to reverse expression of erythroid markers (Fig. 3b,c). To further assess the role of folate metabolism in differentiation of CML cells, we cultured patient-derived CML CD34^+^ cells with low concentration (3U/ml) of erythropoietin, required to sensitise cells to erythroid commitment. As observed with K562 cells, SHIN1 treatment increased levels of GlyA and CD71 in four separate CML CD34^+^ patient samples, while formate supplementation reversed enhanced expression of erythroid markers (Fig. 3d). As AICAR was found to induce differentiation of AML cells^39, 40^, we hypothesised that folate metabolism inhibition would elicit a similar effect on AML cells. To address this, we took advantage of two monocytic AML cell lines, THP1 and MOLM13, which express monocytic surface markers such as CD11b. Increased expression of CD11b correlates with differentiation into the myeloid lineage. Pharmacological inhibition of 1C metabolism resulted in increased expression of CD11b, indicating that SHIN1 treatment induces myeloid differentiation in AML cells, and, therefore, the effect is not specific to K562 cells and erythroid differentiation (Fig. 3d).

Maturation of progenitor cells has been associated with increased oxidative stress and reactive oxygen species (ROS)^43, 44^. Therefore, we measured levels of mitochondrial superoxide following SHIN1 treatment. Surprisingly, SHMT1/2 inhibition resulted in decreased mitochondrial ROS levels (Fig. 3e). Moreover, addition of the ROS scavenger N-acetylcysteine (NAC) did not reverse the enhanced expression of GlyA and CD71 in SHIN1 treated cells (Fig. 3f). These data suggest that initiation of differentiation following inhibition of folate metabolism is not a result of increased ROS production. We next sought to investigate whether AMPK activation was playing a role in differentiation of CML cells. We generated AMPKα1 AMPKα2 double-knockout (DKO) cells (Extended Data Fig. 3e), and measured levels of erythroid markers following incubation with either SHIN1 or AICAR. Surprisingly, there was no significant difference in expression of erythroid markers between SHIN1-treated control and AMPKα DKO cells, indicating that initiation of differentiation is not dependent on AMPK activity (Fig. 3g). Intriguingly, inhibition of folate metabolism suppressed mTORC1 activity in AMPKα DKO cells, measured by decreased phosphorylation of ULK1 on serine-757 and RPS6 (Fig. 3h). The effect of AICAR on mTORC1 activity was reversed in AMPKα DKO cells, suggesting that AMPK signalling is required for AICAR-induced mTORC1 suppression (Fig. 3h).

mTORC1 and AMPK integrate energy signals to balance anabolic and catabolic pathways, including autophagy. When assessing levels of autophagy markers in AMPKα DKO cells, we observed that SHIN1 promoted reduction in the autophagy cargo receptor sequestosome 1 (SQSTM1/p62) and accumulation of the microtubule-associated protein 1 light chain 3B (LC3B-II), indicating induction of the autophagic flux (Fig. 3h). To assess whether autophagy inhibition could reverse the SHIN1-driven differentiation, we utilised two autophagy inhibitors, hydroxychloroquine (HCQ), which blocks the fusion of the autophagosome with the lysosome^45^, and a recently developed inhibitor of ULK1, MRT403^25^. HCQ and MRT403 were sufficient to reverse SHIN1-driven differentiation in control and AMPKα DKO cells (Fig. 3i). Furthermore, SHIN1 treatment did not significantly induce expression of GlyA and CD71 in autophagy deficient, ULK1 KO and autophagy related 7 (ATG7) KO cells (Extended data Fig. 3f,g). Collectively, these data suggest that inhibition of mitochondrial 1C metabolism induces differentiation of CML cells in an AMPK independent manner, while autophagy inhibition reverses such effect.

### Impairment of purine biosynthesis drives differentiation of CML cells through inhibition of mTORC1 signalling

Having established that suppression of mitochondrial folate metabolism drives differentiation and suppression of mTORC1 activity independently of AMPK, we asked whether there is a link between mTORC1 signalling and differentiation. It has been previously described that mTORC1 senses acute changes in purine nucleotide levels, and more specifically adenylates^46, 47^. To determine if this is the case in CML cells, we supplemented hypoxanthine in SHIN1-treated or SHMT2 KO cells and assessed mTORC1 activity. As expected, SHIN1 treatment decreased phosphorylation of the direct mTORC1 kinase target S6 (S6K) and RPS6, while hypoxanthine rescued mTORC1 activity (Fig. 4a and Extended Data Fig. 4a). Importantly, short-term (1 h) supplementation of hypoxanthine was sufficient to reverse mTORC1 signalling, supporting the notion that mTORC1 can acutely sense changes in purine levels. As anticipated, mTORC1 was not reactivated following thymidine supplementation (Fig. 4a and Extended Data Fig. 4a). Furthermore, long-term (24 h) hypoxanthine supplementation repressed AMPK signalling in SHIN1-treated cells, whereas short-term hypoxanthine had a modest effect on AMPK activity (Extended Data Fig. 4b). The pattern of mTORC1 signalling strongly correlated with expression of erythropoiesis markers, with hypoxanthine reversing enhanced levels of GlyA and CD71 (Fig. 4b). In addition to hypoxanthine, adenine treatment restored mTORC1 activity (Fig. 4c and Extended Data Fig. 4c). Long-term guanine supplementation partially reactivated mTORC1, whereas short-term supplementation did not have any effect on mTORC1 signalling. Moreover, only adenine, but not guanine supplementation, repressed increased levels of erythroid maturation markers following 1C metabolism inhibition (Fig. 4d). Similarly, while long-term adenine supplementation fully restored mTORC1 signalling and reversed differentiation in AMPKα DKO, guanine partially reactivated mTORC1, which was not sufficient to reverse differentiation (Fig. 4e,f). To further assess whether mTORC1 inhibition was sufficient to induce differentiation, we treated CML cells with rapamycin and measured levels of erythropoiesis markers. Rapamycin enhanced levels of GlyA and CD71 to the same extent as SHIN1 (Extended Data Fig. 4d). Of note, combination of SHIN1 with rapamycin modestly increased levels of GlyA and CD71 compared to SHIN1 monotherapy, however this did not reach statistical significance (Extended Data Fig. 4d). Ultimately, to confirm the role of mTORC1 activity in CML differentiation, we generated tuberous sclerosis complex 2 (TSC2) KO cells (Fig. 4g). TSC2 is part of the tumour suppressor TSC complex that maintains mTORC1 inactive through the regulation of Ras homologue enriched in brain (Rheb). Additionally, TSC2 has been shown to be required for purine sensing by mTORC1^46^. Intriguingly, loss of TSC2 diminished the effect of SHIN1 on mTORC1 signalling and differentiation in CML cells (Fig. 4g,h). As expected, rapamycin suppressed mTORC1 activity and induced expression of erythroid maturation markers in TSC2 KO cells in line with Fig. S4D. Together, these data uncover that mitochondrial 1C metabolism inhibition mediates differentiation through modulation of the mTORC1 pathway.

**Fig. 4:**
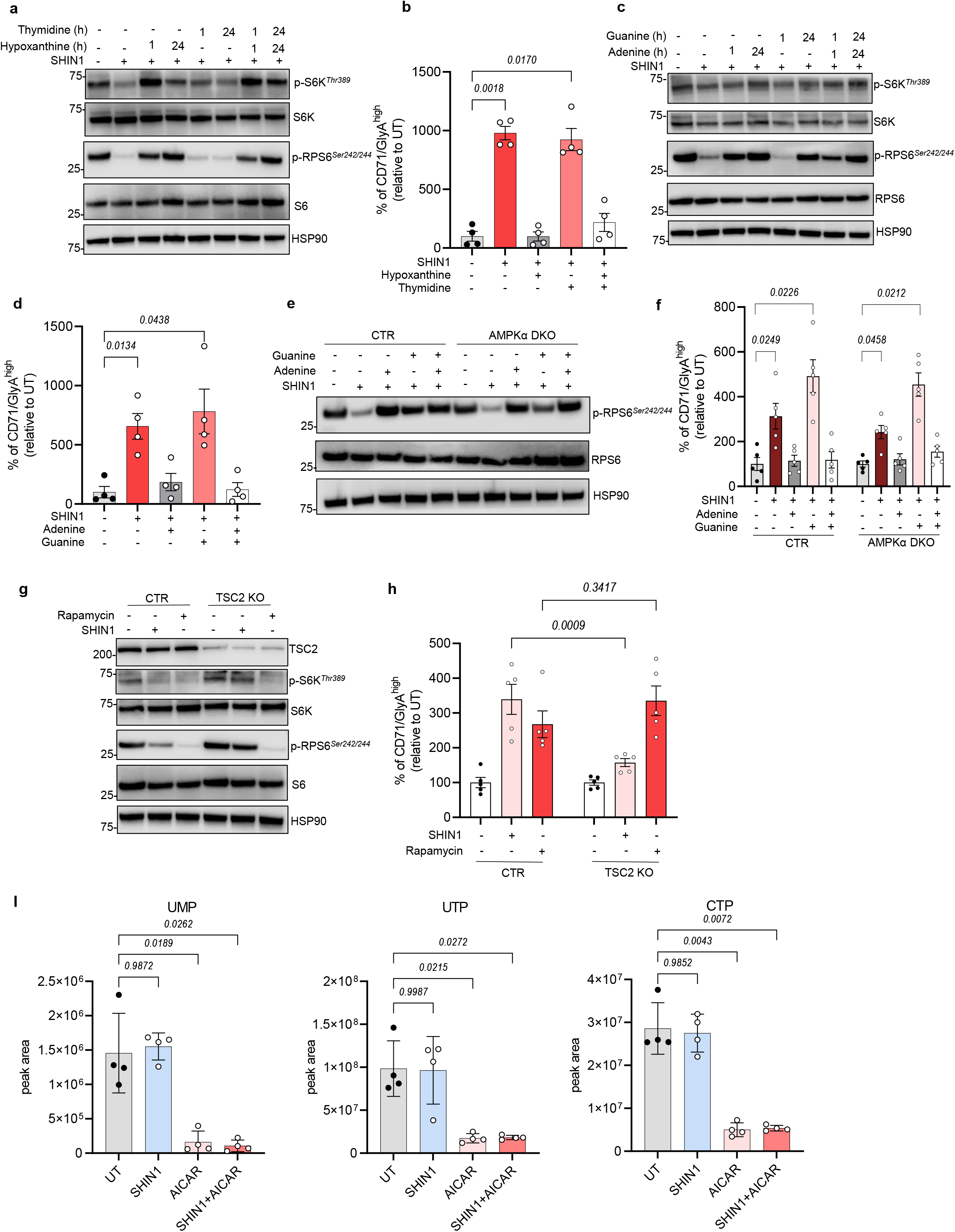
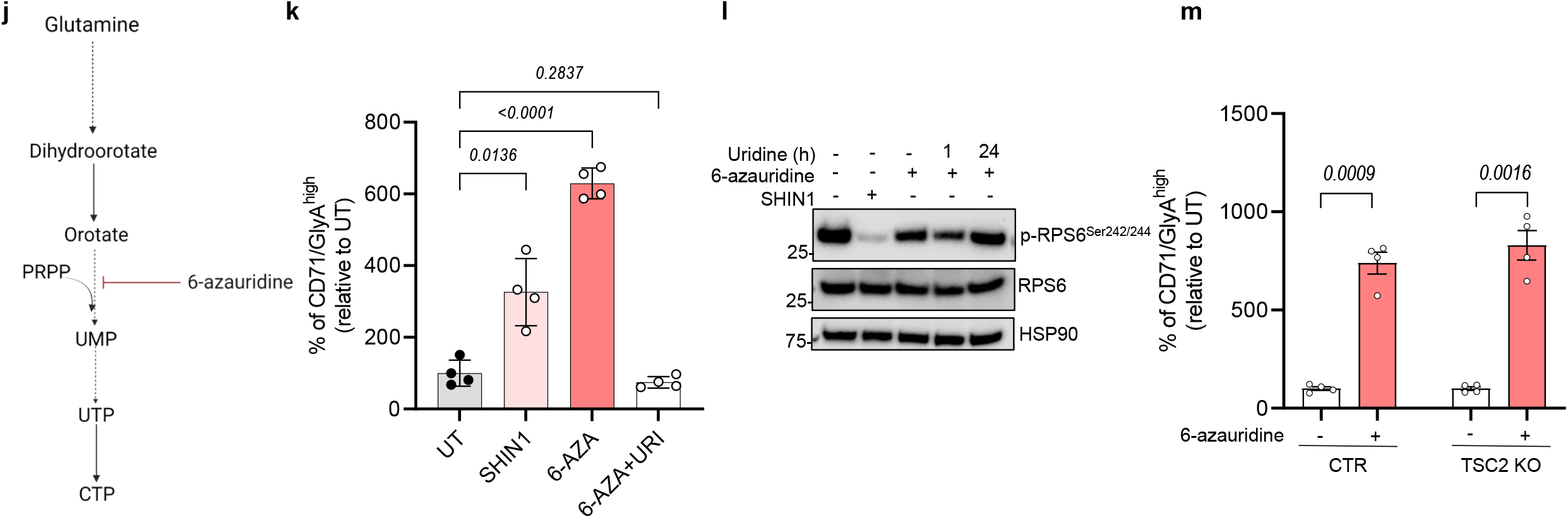
Impairment of nucleotide biosynthesis drives differentiation of CML cells. **a**, Western blot analysis of mTORC1 downstream targets in K562 cells following 24 h treatment with 2.5 μM SHIN1 in the presence or absence of 100 μM hypoxanthine and/or 16 μM thymidine for the duration of the treatment or for the final hour (representative of three independent experiments). **b**, Percentage of CD71/GlyA^high^ K562 cells relative to untreated after 72 h incubation with 2.5 μM SHIN1 with or without the addition of 100 μM hypoxanthine and/or 16 μM thymidine (n=4 independent cultures). **c**, Western blot analysis of mTORC1 targets in K562 cells following 24 h treatment with 2.5 μM SHIN1 in the presence or absence of 30 μM adenine and/or 30 μM guanine for the duration of the treatment or for the final hour (representative of three independent experiments). **d**, Percentage of CD71/GlyA^high^K562 cells relative to untreated after 72 h incubation with 2.5 μM SHIN1 with or without the addition of 30 μM adenine and/or 30 μM guanine (n=4 independent cultures). **e**, Immunoblot analysis of RPS6 phosphorylation in K562 control and AMPKα KO cells incubated with 2.5 μM SHIN1 with or without the addition of 30 μM adenine and/or 30 μM guanine for 24 h (representative of three independent experiments). **f**, Relative percentage of CD71/GlyA^high^ K562 control and AMPKα DKO cells following treatment with 2.5 μM SHIN1 with or without the addition of 30 μM adenine and/or 30 μM guanine (n=4 independent cultures). **g**, Immunoblot analysis of mTORC1 downstream targets in K562 control and TSC2 KO cells incubated with 2.5 μM SHIN1 or 10nM rapamycin for 24 h (representative of three independent experiments). **h**, Relative percentage of CD71/GlyA^high^ K562 control and TSC2 KO cells following treatment with 2.5 μM SHIN1 or 10 nM rapamycin (RAPA) (n=5 independent cultures). **i**, LC-MS measurement of intracellular pyrimidine nucleotides (UMP, UTP, CTP) following 24 h incubation with 1 mM AICAR, 2.5 μM SHIN1, or a combination of both (n=4 independent cultures). **j**, Schematic of 6-azauridine targeting pyrimidine biosynthesis. **k**, Percentage of CD71/GlyA^high^ K562 cells relative to untreated after 72 h incubation with 2.5 μM SHIN1, 5 μM 6-azauridine with or without the addition of 100 μM uridine (URI) (n=4 independent cultures). **l**, Immunoblot analysis of RPS6 phosphorylation in K562 cells following treatment with 2.5 μM SHIN1, 5 μM 6-azauridine in the presence or absence of 100 μM uridine for the duration of the treatment or for the final hour (representative of three independent experiments). **m**, Relative percentage of CD71/GlyA^high^ K562 control and TSC2 KO cells following treatment with 5 μM 6-azauridine (n=4 independent cultures). Data are shown as the mean ±s.e.m. P-values were calculated with a repeated measure one-way ANOVA with Dunnett’s multiple comparisons test (**b,d,f,i,k**), two-way ANOVA with Sidak’s multiple comparisons test (**H**) or two-tailed paired t-test (**m**).

Having determined a link between inhibition of folate metabolism, purine sensing through mTORC1 and differentiation, we sought to further explore the mechanism behind AICAR-induced differentiation. Metabolomic analysis revealed that AICAR treatment significantly decreased the pyrimidine nucleotide pool, while it did not have any significant effect on purine nucleotides (Fig. 4i and Extended Data Fig. 4e). It is also worth noting that SHIN1 did not alter levels of pyrimidine nucleotides (Fig. 4i). These findings are in agreement with previous literature suggesting that AICAR induces differentiation through pyrimidine depletion^40^. To further examine whether impairment of *de novo* pyrimidine synthesis results in differentiation of CML cells, we treated K562 cells with the uridine analogue 6-azauridine. 6-azauridine blocks the conversion of orotate to uridine monophosphate (UMP) (Fig. 4j). Strikingly, impairment of pyrimidine biosynthesis enhanced expression of erythropoiesis markers (Fig. 4k). Uridine supplementation reversed differentiation, supporting that 6-azauridine treatment induced maturation through depletion of pyrimidine nucleotides (Fig. 4k). Moreover, 6-azauridine had a minimal effect on mTORC1 activity, measured by decreased phosphorylation of RPS6, although not to the same extent as SHIN1 (Fig. 4l). Pyrimidine depletion through 6-azauridine treatment mediated erythroid maturation in TSC2-deficient cells (Fig. 4m), suggesting that differentiation following pyrimidine impairment is not dependent on mTORC1 modulation. Overall, these results reveal that impairment of *de novo* nucleotide synthesis results in differentiation of CML cells.

### SHIN1 selectively targets LSCs in combination with standard CML therapy

As previously highlighted, CML is a paradigm for stem cell driven malignancies, as it originates in a single HSC following the generation of the BCR-ABL1 fusion oncoprotein. While the introduction of TKIs such as imatinib has significantly improved response rates of CML patients, disease initiating LSCs are insensitive to TKI therapy^28^. Therefore, having established that folate metabolism is deregulated in LSCs (Fig. 1b), we sought to investigate the combinatory effect of SHIN1 and imatinib treatment on 1C metabolism and survival of primitive CML cells. First, we used stable isotope labelled ^13^C_3_^15^N_1_-serine to determine the effect of imatinib on serine metabolism in K562 cells. Due to SHMT reversibility, serine’s labelling pattern consists of a mixture of m+1, m+3, m+4 isotopologues, reflecting the different recombination events with labelled and unlabelled glycine and 1C units derived from methylene-THF. As expected, SHIN1 treatment increased the abundance of m+4 serine isotopologue, while it dramatically decreased m+1 and m+3 fractions (Fig. 5a and Extended Data Fig. 5a). Additionally, it resulted in decreased abundance of m+3 glycine isotopologue (Fig. 5a). Of note, imatinib did not alter the serine and glycine labelling pattern, suggesting that SHMT activity was maintained (Fig. 5a and Extended Data Fig. 5a). Through generation of glycine, folate metabolism contributes to glutathione (GSH) production, the most abundant antioxidant in the cell. Strikingly, serine incorporation into GSH and its oxidised form (GSSG) persisted in TKI-treated K562 cells, while SHIN1 completely abrogated the contribution of extracellular serine to GSH and GSSG (Fig. 5b and Extended Data Fig. 5b). It should be mentioned that imatinib, similarly to SHIN1, reduced serine labelling into the purine nucleotide pool (Fig. 5c and Extended Data Fig. 5c). Furthermore, serine contribution to glycine, GSH and GSSG was maintained in imatinib-treated primary CML CD34^+^ cells, while SHIN1, and consequently, combination of SHIN1 with imatinib, significantly reduced levels of relevant labelled fractions in GSH and GSSG (Fig. 5d,e and Extended Data Fig. 5d). These data suggest that TKI therapy does not affect the incorporation of serine into GSH, implying a sustained ability for TKI treated CML cells to maintain redox defence.

**Fig. 5:**
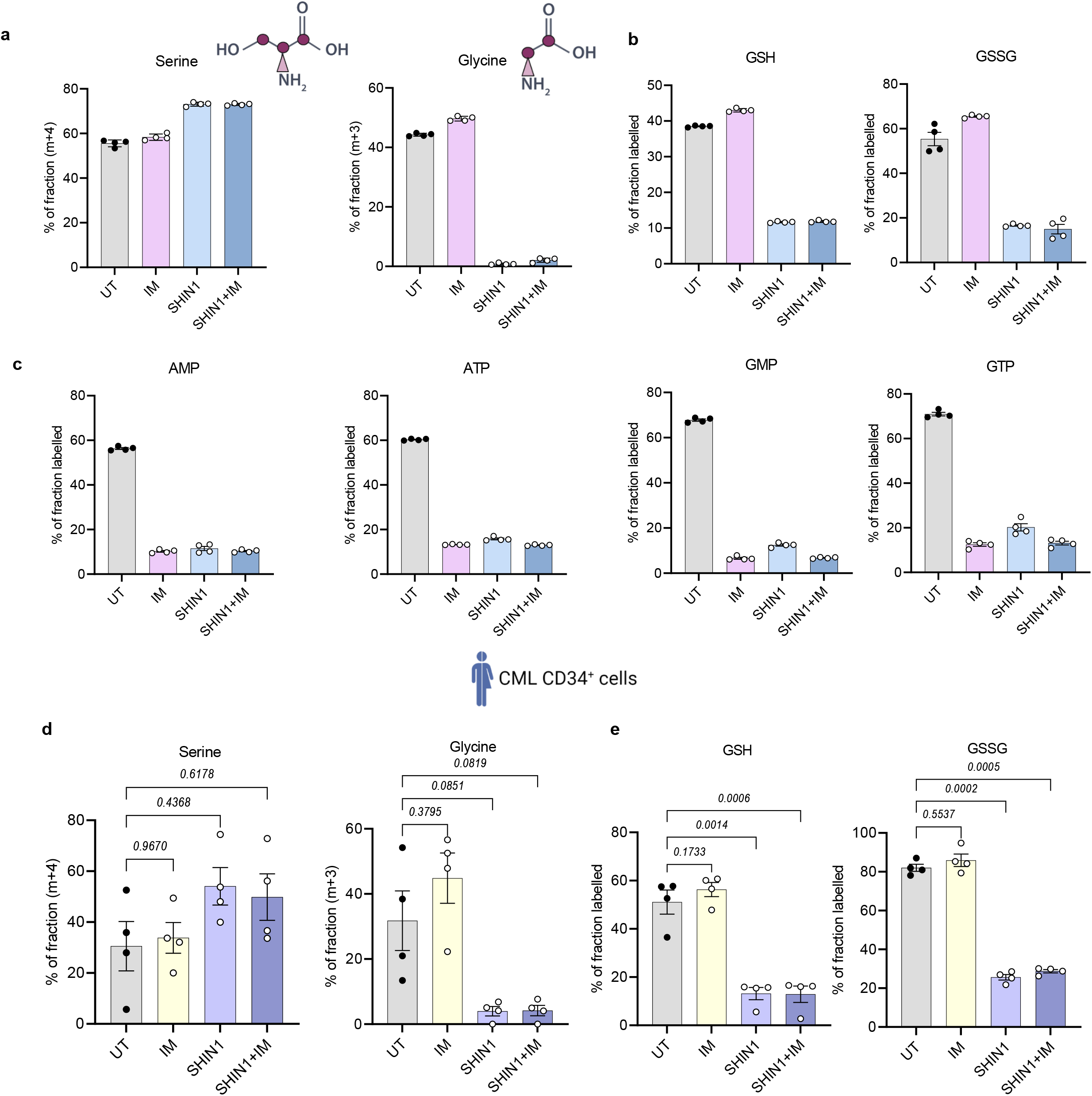
Imatinib maintains serine contribution to glutathione. **a,b,c** The fractional labelling of serine, glycine (**a**), GSH, GSSG (**b**), and purine nucleotides (AMP, ATP, GMP, GTP) (**c**) from K562 cells treated for 24 h with 2 μM imatinib (IM), 2.5 μM SHIN1, or a combination of both in medium containing 140 μM ^13^C_3_^15^N_1_-serine. (n=4 independent wells from individual experiment). **d,e,**Main isotopologue(s) of serine, glycine (**d**) and GSH, GSSG (**e**) from CML CD34^+^ cells treated for 24 h with 2 μM imatinib (IM), 2.5 μM SHIN1, or a combination of both in medium containing 140 μM ^13^C_3_^15^N_1_-serine (n=4 patient samples). Isotopologues are plotted as the percentage of fraction of the sum of all isotopologues. Data are shown as the mean ± s.e.m. P-values were calculated with a repeated measure one-way ANOVA with Dunnett’s multiple comparisons test (**d,e**).

Given the antiproliferative effect observed following mitochondrial folate metabolism inhibition in K562 cells and *in vivo*, we labelled CML CD34^+^ cells with a fluorescent tracer of cell division (CTV) and treated them with imatinib, SHIN1 or a combination of both. Simultaneously, we supplemented SHIN1-treated cells with formate, to assess whether we could recapitulate the findings obtained in K562 cells. SHIN1, alone or in combination with imatinib, impaired the proliferation of primary CML CD34^+^ cells, even to a greater extent than imatinib alone (Fig. 6a,b). Formate supplementation restored proliferation in SHIN1 treated cells (Fig. 6a,b). Furthermore, SHIN1 and imatinib single treatments reduced the colony formation potential of patient-derived CD34^+^ cells (n=7) in colony-forming cell (CFC) assay (Fig. 6c,d). Moreover, SHIN1 sensitised CD34^+^ cells to TKI treatment. Formate was sufficient to reverse the effect of SHIN1 on colony formation potential of primitive CML cells (Fig. 6c,d). In contrast, neither single agent nor combination treatment affected the colony forming potential of normal CD34^+^ cells (Fig. 6e). Encouragingly, using two separate primary CML samples, a combination of folate metabolism inhibition with TKI therapy reduced colony formation in a long-term culture-initiating cell (LTC-IC) assay, the most stringent *in vitro* stem cell assay, suggesting that 1C metabolism is critical for the survival of CML LSCs (Fig. 6f).

**Fig. 6:**
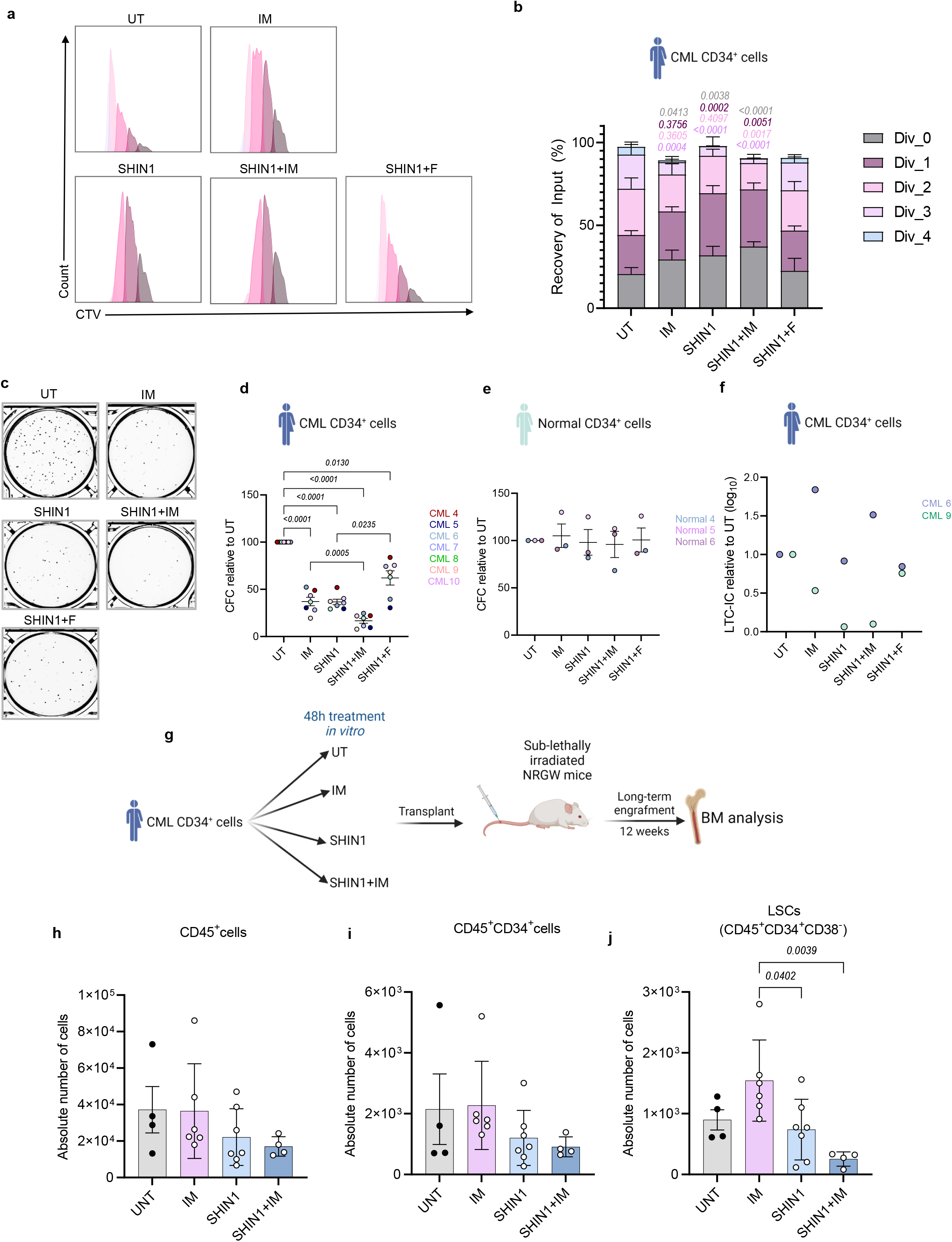
SHIN1 selectively targets LSCs in combination with standard CML therapy. **a**, Representative flow cytometry histograms obtained from cell division tracking of CML CD34^+^ cells using CellTrace Violet (CTV) staining following 72 h of treatment with 2 μM imatinib, 2.5 μM SHIN1 with or without the addition of 1mM formate, or a combination of imatinib and SHIN1. **b**, CTV tracked proliferation of CML CD34^+^ cells as depicted in (**a**), reported as recovery of input (%), where input represents number of input cells (n=4 patient samples). **c,d,**Representative images of colonies (**c**) and relative colony numbers (**d**) after exposure of CML CD34^+^ to treatment as in (**a)** (n=7 patient samples). **e**, Relative colony numbers of normal CD34^+^ cells after treatment as explained in (**a)** (n=3 patient samples). **f**, Relative number of colonies measured by LTC-IC assay after exposure of CML CD34^+^ cells to treatment for 72 h as in (**a-e**) (n=2 patient samples). **g**, Schematic overview of PDX experimental design: CML CD34^+^ cells were treated with 2 μM imatinib, 2.5 μM SHIN1, or a combination of both for 48 h, and transplanted into sub-lethally irradiated female NRGW^41^ mice. After 12 weeks, bone marrow was collected, and engraftment determined by flow cytometry. **h-i**, Number of human CD45^+^ (**h**), CD34^+^ (**i**), and CD34^+^CD38^-^ (**j**) CML cells in bone marrow at experimental endpoint (n=4 mice for untreated group; n=6 mice for imatinib-treated group; n=7 mice for SHIN1-treated group, n=4 mice for combination group). Data are shown as the mean ± s.e.m. P-values are derived from two-way ANOVA with Tukey’s multiple comparisons test (**b**) relative to untreated CML CD34^+^ cells or ordinary one-way ANOVA with Dunnett’s multiple comparisons test (**d,j**).

To further examine the clinical relevance of our findings, we utilised a robust CML xenotransplantation model. Since SHIN1 is not suitable for *in vivo* treatment^33^, we treated patient-derived CML CD34^+^ cells with imatinib, SHIN1 or a combination of both for 48 h, and transplanted them into sub lethally irradiated NRGW (*NOD.Cg-Rag1^tm1Mom^Kit^W-41J^Il2rg^tm1Wjl^*)^41^ mice (Fig. 6g). After 12 weeks, bone marrow cells were collected, and engraftment determined by flow cytometry (Extended Data 6a). We observed a reduction in the percentage of human CD45^+^ leukocytes and total number of human bone marrow CD45^+^ cells in the SHIN1 and imatinib combination group when compared to imatinib alone, although this did not reach statistical significance (Fig. 6h and Extended Data Fig. 6b). Similar results were obtained for the total and relative number of human CD45^+^CD34^+^ cells (Fig. 6i and Extended Data Fig. 6c). However, when examining the most primitive population of human LSCs (CD45^+^CD34^+^CD38^-^), combination of SHIN1 with imatinib conferred a 75% reduction of the absolute number of LSCs compared to imatinib alone (Fig. 6j). A significant decrease in the relative number of human LSCs was also observed (Extended Data Fig. 6d). Notably imatinib increased the total number of LSCs, supporting literature that suggests that TKI therapy enriches for the most primitive population of LSCs (Fig. 6j). Together these data highlight that 1C metabolism inhibition sensitises therapy resistant LSCs to targeted therapy.

## Discussion

Our findings demonstrate an undescribed role for mitochondrial folate metabolism in CML, a malignancy with a well-established cancer stem cell hierarchy. Previous work from our lab highlighted the importance of mitochondrial metabolism, and particularly OXPHOS, in therapy resistant LSCs^29^. Furthermore, it has been reported that increased serine catabolism to formate is a hallmark of oxidative cancers^8^. However, while targeting folate metabolism is an active area of research in cancer, the role of 1C metabolism in CML LSCs is unexplored. We show that folate metabolism is deregulated in human stem and progenitor cells. This deregulation is mainly driven by the differential expression of three mitochondrial-specific enzymes, ALDH1L2, SHMT2 and MTHFD2. Moreover, we found that mitochondrial folate metabolism, primarily through its contribution to purine biosynthesis, is essential for CML cell proliferation and tumour formation in mice. Specifically, we demonstrate that inhibition of mitochondrial 1C metabolism through loss of SHMT2 inhibits tumour formation capacity of CML cells. This is in contrast to work from Ducker *et al*.^5^ which showed that HCT-116 colon cancer cells, engineered with mitochondrial 1C enzyme deletions, form xenograft tumours by generating cytosolic 1C units from serine via the enzyme SHMT1.

Additionally, our work demonstrates that depletion of 1C units and purine nucleotides results in differentiation of CML K562 cells and patient-derived progenitor cells. We show that the differentiation is not a consequence of increased ROS, one of the main drivers of differentiation in HSCs. We also conclude that initiation of differentiation does not depend on AMPK activity but is linked to purine sensing through mTORC1 signalling. Recent work has described that impairment of purine biosynthesis leads to inactivation of mTORC1^46, 47^. Furthermore, it has also been reported that mTORC1 promotes serine catabolism to formate and purine biosynthesis through induction of the activating transcription factor 4 (ATF4)^48^, supporting the existence of a positive feedback loop between folate metabolism, *de novo* purine synthesis and mTORC1 signalling. Here, we observe that while pharmacological inhibition of 1C metabolism leads to suppression of mTORC1 activity, this can be rescued either by reconstitution of 1C units or purine nucleotides, preferably adenine. These findings are consistent with the study of Hoxhaj *et al*.^46^ which demonstrated that mTORC1 acutely senses changes in adenylates rather than guanylates. Furthermore, our data support that reactivation of mTORC1 correlates with suppression of differentiation. Ultimately, we found that purine depletion fails to suppress mTORC1 activity and induce erythroid maturation in TSC2-deficient cells, supporting that purine sensing through the mTORC1 network mediates differentiation of CML cells. Additionally, we described that pyrimidine depletion initiates differentiation of CML cells, while it only modestly affects mTORC1 activity. These data suggest that impairment of pyrimidine biosynthesis induces differentiation through a separate mechanism, independently of mTORC1 signalling. Indeed, in agreement with our data, it has been recently suggested that pyrimidine depletion alters cell fate of AML cells through DNA replication stress^49^

In terms of clinical relevance, we found that folate metabolism inhibition synergises with TKI treatment to target therapy insensitive LSCs. Resistance of LSCs to standard therapy has been associated with their undifferentiated and quiescent state^50, 51^. Thus, we propose that LSCs upregulate folate metabolism to drive leukaemogenesis, without significantly deregulating the normal differentiation process. Intriguingly, Dohla *et al*.^52^ uncovered that in stem-like human mammary epithelial cells, purine synthesis was upregulated in daughter cells that were destined to maintain stemness. We suggest that inhibition of folate metabolism through impairment of purine biosynthesis can prime LSCs out of quiescence and thereby sensitise them to TKI therapy, which preferentially targets more differentiated cells. Notably, TKI therapy was found to maintain serine incorporation to GSH and GSSG, suggesting that the synergistic effect with folate metabolism inhibition might be related to impeded antioxidant defence. Serine metabolism contributes to the maintenance of the GSH pool in two ways. Firstly, serine-derived glycine combines with cysteine and glutamate to form GSH. Secondly, serine catabolism contributes to generation of mitochondrial NADPH^53, 54^, which is consumed during the reduction of GSSG to GSH. Whether alterations of the GSH pool sensitises cells to TKI therapy requires further investigation.

Lastly, it has been shown that suppression of MTHFD2 leads to differentiation of AML cells. Pikman *et al*.^55^ demonstrated that formate supplementation did not restore proliferation following suppression of MTHFD2, implying that the antiproliferative effect of MTHFD2 ablation was not a result of 1C unit depletion. In the present study, we describe that folate metabolism inhibition initiates differentiation of AML cells, and this can be reversed by addition of formate. Moreover, together these data indicate that mitochondrial 1C metabolism is a targetable vulnerability for myeloid leukaemias and suggest that folate metabolism-driven differentiation may have a broader application to related malignancies of stem cell origin. Given that *SHMT2* and *MTHFD2* are overexpressed in cancer, mitochondrial 1C metabolism inhibitors might be more selective than cytosolic antifolates like methotrexate. Methotrexate inhibits the activity of dihydrofolate reductase (DHFR), but has affinity for folate-dependent enzymes such as thymidylate synthase (TYMS) and 5-aminoimidazole-4-carboxyamide ribonucleotide formyltransferase^56, 57^. It has been shown that DHFR and TYMS are expressed in many proliferating tissues, in which the adverse effects of MTX are observed^7^. In contrast, we demonstrate that SHIN1 does not have any effect in normal CD34^+^ cells, supporting the concept that inhibiting mitochondrial 1C metabolism may be more selective and have less adverse side effects.

## Methods

### Patient derived samples and approval

Primary CML samples were obtained from individuals in chronic phase CML at the time of diagnosis and were product of leukapheresis. Normal samples were (i) surplus cells collected from femoral-head bone marrow, surgically removed from patients undergoing hip replacement or (ii) leukapheresis products from individuals with non-myeloid Philadelphia negative haematological disorders. CD34^+^ cells were isolated using the CD34 MicroBead Kit or CliniMACS (catalogue no. 130-100-454; Miltenyi Biotec). Patients gave informed consent in accordance with the Declaration of Helsinki and with approval of the National Health Service Greater Glasgow Institutional Review Board and Clyde Institutional Review Board. Ethical approval has been granted by the West of Scotland Research Ethics Service (REC reference: 15/WS/0077 and 10/S0704/60). Patient information is available in Supplementary Table 1.

### Cell culture

CML and AML cell lines were cultured in RPMI 1640 medium (catalogue no. 11875093; Thermo Fisher Scientific) supplemented with 100 IU/mL penicillin/streptomycin, 2 mM L-glutamine (catalogue no. A2916801; Thermo Fisher Scientific), and 10% (v/v) fetal bovine serum (FBS, catalogue no. 10100147). SHMT2 KO cells were cultured with 1% hypoxanthine/thymidine supplement (HT, catalogue no. 11067030; Thermo Fisher Scientific). HT supplement was removed prior to any experimental procedures. Primary samples were cultured in serum-free medium (SFM) consisting of IMDM (catalogue no. 12440061; Thermo Fisher Scientific) and supplemented with 10% bovine serum albumin (BSA), insulin, and transferrin (BIT, catalogue no. 09500; STEMCELL Technologies), 40 μg/ml low-density lipoprotein (catalogue no. L4646; Sigma-Aldrich), 0.1 mM 2-mercaptoethanol (catalogue no. 31350-010; Thermo Fisher Scientific) and 100 IU/mL penicillin-streptomycin (catalogue no. 15140122; Thermo Fisher Scientific). Cells were also supplemented with a physiological growth factor cocktail consisting of 0.2 ng/ml stem cell factor (SCF, catalogue no. 573902; BioLegend), granulocyte macrophage colony-stimulating factor (GM-CSF, catalogue no. 578602; BioLegend), 1 ng/ml interleukin-6 (IL6, catalogue no. 570802; BioLegend), macrophage inflammatory protein (MIPα, catalogue no. 300-08; PeproTech) and 0.05 ng/ml leukemia inhibitory factor (LIF, catalogue no. 300-05; PeproTech). Both cell lines and primary cells were maintained at 5% CO_2_ and 37°C. Cell lines were routinely tested for mycoplasma. Cell numbers were monitored using the CASY automated cell counter (Roche).

Where indicated, the following compounds were added to the medium: 2.5 μM SHIN1 (catalogue no. HY-112066; Cambridge Biosciences), 1 mM AICAR (catalogue no. A8184-APE; ApexBio), 10 nM rapamycin (catalogue no. R-5000; LC Laboratories), 5 μM 6-azauridine (catalogue no. A1888; Sigma-Aldrich), 5 μM HCQ (catalogue no. 10646271; Fischer Scientific), 3 μM MRT403 (LifeArc), 10 nM rapamycin (catalogue no. R-5000, LC Laboratories), and 2 μM imatinib mesylate (catalogue no. I-5508; LC Laboratories). For formate supplementation experiments, sodium formate (catalogue no. 141-53-7; Sigma-Aldrich) was added at a concentration of 1 mM. Hypoxanthine (catalogue no. A11481; Thermo Fisher Scientific) and thymidine (catalogue no. A11493.06; Thermo Fisher Scientific), were used in a concentration of 100 μM and 16 μM respectively. For purine nucleotide supplementation, cells were cultured in 30 μM adenine (catalogue no. A2786; Sigma-Aldrich) or/and 30 μM guanine (catalogue no. G6779; Sigma-Aldrich). For pyrimidine rescue experiments, uridine (A15227; Thermo Fisher Scientific) was added at a concentration of 100 μM.

### Growth curves and proliferation rates

K562 cells were seeded at a density of 50,000 cells/ml in a 12-well plate. Indicated treatments were added to separate wells (three wells per condition). Cell counts were obtained at 24h intervals using the CASY automated cell counter (Roche). The proliferation rate was determined based on the following formula:

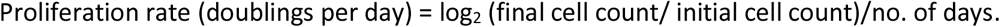

### Cell cycle analysis

K562 cells were cultured at 100,000 cells/ml in the presence of the indicated treatments. Following a 72 hours (h) incubation, cells were fixed in cold 70% ethanol while vortexed and then incubated at −20°C for at least 2 h. Cells were then stained with propidium iodide (PI, catalogue no. P1304MP; Thermo Fisher Scientific) in the presence of 100 μg/ml of RNase A (QIAGEN) for 20 min and analysed using the FaCSVerse flow cytometer (BD Biosciences). Analysis of cell cycle stages using PI mean fluorescence intensity (MFI) was performed with FlowJo V10.7.

### Generation of knockout and stable cell lines

To generate CRISPR–Cas9-mediated gene knockouts in CML cells, we used guide sequences as described in Supplementary Table 2. Synthesis of selected guides was conducted by Integrated DNA Technologies. Oligonucleotides were annealed and cloned in BsmBI–digested lentiCRISPRv.2-puro plasmid (catalogue no. 52961; Addgene). Lentiviral particles for cloned lentiCRISPRv.2 were generated by transfecting human embryonic kidney (HEK) 293FT cells with pCMV-VSV-G envelope and psPAX2 packaging vectors using the calcium phosphate method ^58^. Viral supernatant was harvested after 48 h and transferred onto target CML cells. Cells were then selected using 3 μM/ml puromycin and presence of guides was validated by immunoblotting. K562 ULK1 KO and K562 ATG7 KO cells were previously generated as described by Ianniciello *et al*.^25^.

### Western blot analysis

Cells from a six-well plate were rinsed with PBS and lysed into RIPA buffer (catalogue no. 89900; Thermo Fisher Scientific) containing cOmplete Protease (Roche) and PhosSTOP phosphatase (Roche) inhibitors. Cell lysates were cleared by centrifugation at 16,000g for 15 min at 4°C. Protein concentration was quantified by bicinchoninic acid assay (Thermo Fisher Scientific). Lysates were resolved in 4 to 12% SDS–polyacrylamide gel electrophoresis gels and transferred to PVDF membranes. Blocking was performed for 1 h in 2% BSA (in Tris-buffered saline, 0.01% Tween (TBS-T)) followed by incubation overnight with indicated primary antibodies (Supplementary Table 3) at 4°C. The next day, membranes were incubated with the appropriate horseradish peroxidase conjugated secondary antibodies (Supplementary Table 3) at room temperature for 1 h. Bands were detected by ECL reagent (catalogue no. RPN2235; Cytiva) and developed using the Odyssey FC imaging system.

### Erythroid maturation analysis

To evaluate erythroid differentiation expression of cell surface markers GlyA and CD71 was assessed. K562 cells were cultured at 100,000 cells/ml in the presence of the indicated treatments. CML CD34^+^ cells were cultured at 100,000 cells/ml in SFM medium with the addition of EPO (3 U/ml, catalogue no. 100-64; PeproTech). Following a 72h incubation, cells were stained with fluorophore-conjugated antibodies GlyA-PE (catalogue no. 349106; Biolegend) and CD71-APC (catalogue no. 33410; Biolegend) for 20 min and analysed using the BD FacsVerse flow cytometer.

### Myeloid maturation analysis

Expression of the monocytic cell surface marker CD11b was assessed in AML cell lines. Following 72 h incubation with the indicated treatments, THP1 and MOLM13 were stained with CD11b-PE (catalogue no. 301306; Biolegend) for 20 min and analysed by flow cytometry.

### Mitochondrial ROS measurements

K562 cells (25,000 cells/ml) were treated with the indicated treatments for 72 h and stained with 5μM MITOSOX (catalogue no. M36008; Thermo Fisher Scientific) in 1 ml of phosphate buffered saline (PBS) solution. Cells were then incubated for 30 min at 37°C and analysed by flow cytometry.

### Cellular division tracking

CML CD34^+^ cells were stained with 1 μM CellTrace Violet (catalogue no. C34557; Thermo Fisher Scientific) for 30 min at 37 °C. The reaction was quenched by adding cell culture medium containing 10% FBS. Cells were then suspended in SFM and treated with the indicated treatments. After 72 h, CellTrace Violet staining was assessed by the BD FACSVerse flow cytometer. Analysis was conducted using FloJo 10.7. Each division was gated using the geometric MFI to determine the divisions, with each subsequent division having half the fluorescent intensity of the previous division. Data were when normalised to cell counts collected after 72 h and reported as recovery of input (%), relative to the number of input cells.

### Steady-state metabolic profiling

For steady-state metabolite measurements, CML cells were seeded in different experimental conditions in standard RPM1 1460 medium one day prior to metabolite extraction. On the day of the metabolite extraction, cells were counted using the CASY automated cell counter. Cell number was multiplied by cell size (peak volume) and this number was used to calculate the volume of extraction solvent. Cells were washed twice with ice-cold PBS with pulse centrifugation (12,000g, 15 sec, 4°C) before a 5 min incubation step with extraction buffer (50:30:20, v/v/v acetonitrile/methanol/water). Cells were then centrifuged at 16,000g for 10 min at 4°C, and the supernatant was transferred to liquid chromatography-mass spectrometry (LC-MS) glass vials and kept at −80°C until measurements were performed.

Samples were analysed using a Thermo Ultimate 3000 High pressure liquid chromatography (HPLC) system (Thermo Scientific) coupled to a Q Exactive Orbitrap mass spectrometer (Thermo Scientific). The HPLC system consisted of a ZIC-pHILIC column (SeQuant, 150 × 2.1 mm, 5μm, Merck KGaA), with a ZIC-pHILIC guard column (SeQuant, 20 × 2.1 mm) and an initial mobile phase of 20% 20 mM ammonium carbonate, pH 9.4, and 80% acetonitrile. For the analysis, 5 μl of metabolite extracts were injected into the system, and metabolites were separated over a 15 min mobile phase gradient decreasing the acetonitrile content to 20%, at a flow rate of 200 μl/min and a column temperature of 45°C. Metabolites were then detected using the mass spectrometer operating in polarity switching mode. All metabolites were detected across a mass range of 75 – 1000 m/z at a resolution of 35,000 (at 200 m/z), and with a mass accuracy below 5 ppm. Data were acquired using Thermo Xcalibur software. The peak areas of different metabolites were determined using the TraceFinder 4.1 software (Thermo Fischer Scientific). Metabolites were identified by accurate mass of the singly charged ion and by known retention times on the pHILIC column.

### Stable isotope enrichment analysis using ^13^C_3_^15^N_1_-serine

For tracing experiments, K562 cells and primary CML cells were cultured in Plasmax, a physiological cell culture medium^59^, containing 140 μM ^13^C_3_^15^N_1_-serine (catalogue no. CNLM-474-H-PK; Cambridge Isotope Laboratories). K562 cells were cultured in Plasmax supplemented with 10% dialysed FBS (catalogue no. A3382001; Thermo Fisher Scientific). Primary CML cells were cultured in Plasmax supplemented with physiological growth factors as described above (see section **Cell culture**), however instead of BIT, medium was supplemented with 1 mg/ml Albumax II (catalogue no. 11021037; Thermo Fisher Scientific), 10 μg/ml insulin (catalogue no. I9278; Sigma-Aldrich), and 7.5 μg/mL transferrin (catalogue no. T4132; Sigma-Aldrich). Cell culture, harvest, metabolite extraction, analysis by LC-MS was performed as described for steady state metabolite profiling (see above). The ^13^C^15^N labelling pattern was determined by quantifying peak areas for the accurate mass of all isotopologues of each metabolite. Labelled isotopologues were defined as mass + n (m+n), where n = number of ^13^C or ^15^N incorporated. Fig.s include clarification whether (m+n) consists of only ^13^C isotopologues, or a combination of ^13^C or ^15^N. We calculated the fractional enrichment, which describes the relative contribution of stable isotopes to the total area of each compound.

### Quantitative analysis of extracellular metabolites

Quantity of extracellular glucose and lactate was determined using the YSI 2900 biochemistry analyser as per manufacturer instructions. The concentration of each metabolite was normalised to cell number and the rate of uptake or secretion per hour was calculated relative to medium only sample. The exchange rate for each metabolite was calculated using the formula below:

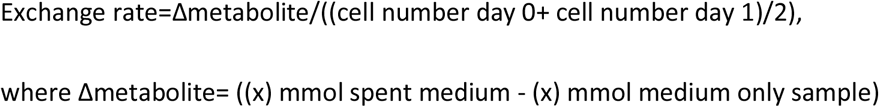

### GC-MS formate analysis

Formate quantification using gas chromatography-mass spectrometry (GC-MS) was performed as previously described by Meiser *et al*.^34^. Media samples from CML CD34^+^ cells (40 μL) were added in 20 μl of 50 μM internal standard sodium ^13^C,^2^H-formate (m+2), 10 μl of 1 M sodium hydroxide, 5 μl of benzyl alcohol and 50 μl of pyridine. While vortexing, derivatisation of formate was commenced by the addition of 20 μl of methyl chloroformate. 200 μl of water and 100 μl of methyl tertiary butyl ether were then added, and the samples were vortexed for 20 sec and centrifuged at 10,000g for 5 min. The resulting apolar phase containing the formate derivative (benzyl formate) was transferred to a GC-vial and capped. Formate standards and blank samples (water) were prepared in the same manner and analysed with the experimental samples. Derivatized formate samples were analysed with an Agilent 7890B GC system coupled to a 7000 triple quadrupole MS system^34^. MassHunter Quantitative analysis software (Agilent Technologies) was used to extract and process the peak areas for m+0, m+1, m+2 formate. After correction for background signals, quantification was performed by comparing the peak area of formate (m/z of 136) and of m+1 formate (m/z of 137) against that of m+2-formate (m/z of 138).

### Measurement of extracellular acidification rate

ECAR of CML cells was measured using the Seahorse XF96 flux analyzer (Seahorse Bioscience). The XF96 well plate was coated with 22.6 μg/ml Cell-Tak solution (catalogue number. 354240; Corning) and left for at least 30 min at room temperature. Following a 24 h incubation with indicated treatments, K562 cells were suspended in XF Assay Medium (catalogue no. 100965–000; Seahorse Bioscience) supplemented with 2 mM glutamine and seeded at a concentration of 70,000-100,000 cells/well in the pre-coated XF96 well plate. Measurement of ECAR was performed at baseline and after sequential injections of glucose (11 mM), oligomycin (0.7 μM), and 2-deoxyglucose (5 mM). Glycolysis and glycolytic capacity relative to untreated or vector control were calculated according to manufacturer’s instructions.

### Computational analysis

Affymetrix microarray dataset (E-MTAB-2581) was analysed using Bioconductor. Packages *oligo* and *arrayQualityMetrics* were used for quality control and pre-processing, while *limma* was used for differential gene expression analysis. Statistical significance was determined using the Benjamini-Hochberg method. Adjusted p-values < 0.05 and log_2_FC ≥ abs (0.5) were considered to be significant. GSEA (version 4.1.0) analysis was conducted on pre-ranked lists (ranked by pi score calculated by multiplying Log_2_ FC by –Log_10_ (adjusted p-value)).

### Colony-forming cell assay

Primary CD34^+^ cells were cultured in SFM in the presence of indicated treatments. After 72 h, 3,000 cells per condition were seeded to methylcellulose based medium (catalogue no. 04034; STEMCELL Technologies) in duplicate, and colonies were counted after 14 days.

### Human long-term culture initiating cell assay

Irradiated M2-10B4 and S1/S1 genetically engineered mouse cell lines, expressing human cytokines, were seeded (70,000 cells each) in medium for long-term culture of human cells (catalogue no. 05150; STEMCELL Technologies) supplemented with hydrocortisone in collagen-coated plates. The following day, 5,000 CML CD34^+^ cells (pre-treat for 72 h) were plated on top of the feeder cell layer and cultured for additional 5 weeks with a weekly refresh of half of the media. Cells were then harvested and assessed by CFC assay

### Tumour xenograft growth

Animal experiments were performed according to all applicable laws and regulations with ethical approval from the University of Glasgow Animal Welfare and Ethical Review Board (AWERB) under Home Office License PPL 60/4492. KCL22 control or SHMT2 KO cells (4,000,000 cells per mouse) were transplanted via tail vein injection into female NRGW^41^ mice aged 8-12 weeks. Mice were euthanized according to institutional guidelines before reaching the threshold maximum tumour burden. Tumour size was recorded using a calibre, and tumour burden was calculated using the formula (volume = 0.5 × (length × width^2^)).

### Mouse engraftment using patient-derived xenografts

To study the *in vivo* engraftment of patient derived CML CD34^+^ cells, CML CD34^+^ cells were cultured in SFM in the presence with appropriate treatment for 48 h. Cells (1,000,000 cells per mouse) were then harvested, washed and transplanted into sub-lethally irradiated (2.0 Gy) female NRGW^41^ mice aged 8-12 weeks. Following 12 weeks, bone marrow was extracted from the hip, tibia, and femur of each mouse. Cells were stained with anti-mouse CD45-APC-Cy7 (catalogue no. 557659; BD Biosciences), anti–human CD45-FITC (catalogue no. 555482; BD Biosciences), anti–human CD34-APC (catalogue no. 555824; BD Biosciences) and anti– human CD38-PerCP (catalogue no.303520; Biolegend) antibodies and analysed by flow cytometry as described above.

### Statistics and reproducibility

Statistical analyses were performed using Prism 9 (Graphpad) software. The number of biological replicates and applied statistical analysis are indicated in the Fig. legends. Statistical significance for parametric data was determined by paired or unpaired two-tailed t-tests. One-way analysis of variance (ANOVA) was used to assess statistical significance between three or more groups with one experimental parameter, while two-way ANOVA was used to determine significance between multiple groups with two experimental parameters. Nonparametric data were analysed using two-sided Mann–Whitney tests. Kruskal-Wallis test was used to assess statistical significance between three or more groups with one experimental parameter, when the data were not normally distributed. For *in vivo* work, mice were randomly allocated to control or experimental conditions. Tumour-free survival was monitored by Kaplan-Meier analysis, and p-values were calculated using log-rank (Mantel-Cox) test. The investigators were not blinded during data collection or analyses.

## Acknowledgments

We would like to thank all members of the Helgason and Vazquez laboratory for input and advice. We thank all patients and healthy donors who donated samples and the National Health Service (NHS) Greater Glasgow and Clyde Biorepository and A. Hair for sample processing. We thank the Core Services and Advanced Technologies at the Cancer Research UK Beatson Institute (C596/A17196; A31287). We thank S. Tardito for providing the components of Plasmax culture medium. We thank C. J. Eaves for providing NRGW^41^ mice and the Cancer Research UK Beatson Institute mouse facility staff for housing of mice and help with xenograft experiments. We thank LifeArc for providing the ULK1 inhibitor, MRT403. MRT403 will be made available to academic researchers by LifeArc upon reasonable request and after completion of a material transfer agreement. We thank J. Jones for reading and editing the manuscript. Schematic graphs in Figs. 1a, 2a, 4h, 5a,b and 6g were generated using BioRender (https://biorender.com).

This work was supported by Cancer Research UK Glasgow Centre (A25142), Cancer Research UK (C57352/A29754), The Kay Kendall Leukemia Fund (KKL1069), Blood Cancer UK (formerly Bloodwise; Ref 18006), The Howat Foundation, Tenovus Scotland and Friends of Paul O’Gorman Leukaemia Research Centre (all to G.V.H.); NHSGGC Endowment Fellowship (GN21ON384) to M.M.Z. and Cancer Research UK to D.S.^2^ (A17196 and A31287) and A.V. (A17196 and A21140). M.M.Z. was supported by a Cancer Research UK PhD scholarship (A25142).

## Author contribution

M.M.Z., A.V. and G.V.H. conceived and designed the study. M.M.Z. performed all experiments and analysed data. K.M.R assisted with stable isotope tracing, primary CD34^+^ CFC experiments and contributed to data interpretation. D.S.^1^ performed microarray dataset analysis and assisted with data interpretation. A.D. and A.I. provided material support and contributed to data interpretation. K.D. assisted with *in vivo* experiments. M.C. provided material support. D.S.^2^ performed the mass spectrometry. A.V. performed the GC-MS data analysis for the formate levels. M.M.Z. and G.V.H. wrote the manuscript and all other authors reviewed it.

## Figure Legends

**Extended data Fig.1: Mitochondrial serine catabolism is required for CML cell proliferation. a,b,** Gene set enrichment analysis of significantly deregulated genes in CML LSCs compared to normal HSCs (CD34^+^CD38^-^) (E-MTAB-2581). NES, normalised enrichment score; FDR, false discovery rate. **c**, Expression of folate metabolism associated gene transcripts plotted in ascending order relative to HSCs. **d**, Formate concentration in media from normal and CML CD34^+^ cells following 48 h culture (n=3 patient samples). **e**, Immunoblot analysis of SHMT2 levels in parental K562 cells as well as K562 expressing an empty vector (CTR) or SHMT2 knockout (KO) cells. **f,g,** Growth (**f**) and proliferation rate (**g**) of K562 CTR and SHMT2 KO cells with or without the addition of 1 mM formate (F), 100 μM hypoxanthine (H), 16 μM thymidine (T) for 72 h (n=3 independent cultures). **h**, Percentage of cell cycle phases of K562 CTR and SHMT2 KO with the addition of treatments as in (**f,g)** (n=3 independent cultures). Data are shown as the mean ± s.e.m. P-values were calculated using unpaired two-tailed t-test with Welch’s correction (**d**), a repeated measure one-way ANOVA with Dunnett’s multiple comparisons test (**g**) or two-way ANOVA with Tukey’s multiple comparison’s test (**h**) relative to control K562 cells.

**Extended data Fig.2: Inhibition of mitochondrial serine catabolism disrupts *de novo* purine synthesis and glycolysis in CML cells. a-d,** LC-MS measurement of intracellular glycine (**a**), ribose-5-phosphate (R5P) (**b**), purine biosynthetic intermediate and nucleotides (**c**) and glycolysis related metabolites (**d**) in K562 CTR and SHMT2 KO cells with or without the addition of 1 mM of formate for 24 h (n=4 independent cultures). **e**, Glucose uptake and lactate secretion in K562 CTR and SHMT2 KO in the presence or absence the addition of 1 mM of formate for 24 h (n=5 independent cultures). **f,g,** Representative extracellular acidification rate (ECAR) profile (**f**), relative glycolysis and glycolytic capacity (**g**) as measured by the Seahorse XF analyser of K562 CTR and SHMT2 KO cells cultured with or without the supplementation of 1 mM of formate (n=4 independent cultures). Data are shown as the mean ± s.e.m. P-values are derived from a repeated measure one-way ANOVA with Dunnett’s multiple comparisons test (**a-e**), ordinary one-way ANOVA with Dunnett’s multiple comparisons test for glycolysis and Kruskal-Wallis test with Dunn’s multiple comparisons test for glycolytic capacity (**g**).

**Extended data Fig.3: AML cells are sensitive to mitochondrial 1C metabolism-induced maturation, while autophagy deficiency prevents differentiation. a,** Immunoblot analysis to assess AMPK and mTORC1 signalling in K562 control and SHMT2 KO cells following 24 h incubation with 1mM AICAR or 1 mM formate (representative of three independent experiments). **b**, Change in pellet colour of K562 cells following SHIN1 treatment or SHMT2 KO. **c**, Percentage of CD71/GlyA^high^ cells relative to control following loss of SHMT2 (n=2 independent cultures). **d**, Quantification of CD11b MFI of THP1 and MOLM13 cells relative to untreated following 72 h incubation with 2.5 μM SHIN1 with or without the supplementation of 1 mM formate (n=5 independent cultures). **e**, Western blot analysis of AMPKα levels and phosphorylation in K562 control and AMPKα DKO cells in the presence of 1 mM AICAR or 2.5 μM SHIN1 for 24 h. **f**, Immunoblot analysis of ULK1 and ATG7 levels in K562 control, ULK1 KO and ATG7 KO cells. **g**, Relative percentage of CD71/GlyA^high^ K562 control, ULK1 KO and ATG7 KO cells treated with 2.5 μM SHIN1 for 72 h. (n=3 independent cultures). Data are presented as the mean ± s.e.m. P-values were calculated using a repeated measure one-way ANOVA with Dunnett’s multiple comparisons test (**d,g**).

**Extended data Fig.4: Role of mTORC1 in nucleotide sensing and differentiation. a, b** Western blot analysis of mTORC1 targets (**a**) and AMPKα phosphorylation (**b**) in K562 control and SHMT2 KO cells in the presence or absence of 100 μM hypoxanthine and/or 16 μM thymidine for the duration of the treatment (24 h) or for the final hour (representative of three independent experiments). **c**, Western blot analysis of mTORC1 targets in K562 control and SHMT2 KO cells with or without the addition of 30 μM adenine and/or 30 μM guanine for the duration of the treatment (24 h) or for the final hour (representative of three independent experiments). **d**, Percentage of CD71/GlyA^high^ K562 cells relative to untreated after 72 h incubation with 10 nM rapamycin (RAPA), 2.5 μM SHIN1, or a combination of both (n=4 independent cultures). **e**, LC-MS measurement of intracellular purine nucleotides (AMP, ADP, ATP, GMP, GDP, GTP) following 24 h incubation with 1 mM AICAR, 2.5 μM SHIN1, or a combination of both (n=4 independent cultures). Data are presented as the mean ± s.e.m. P-values were calculated using repeated two-way ANOVA with Sidak’s multiple comparisons test (**d,g**).

**Extended data Fig.5: Imatinib maintains serine contribution to glutathione. a,b,c** Mass isotopologue distribution in serine, glycine (**a**), GSH, GSSH (**b**) and purine nucleotides (**c**) from K562 cells treated for 24 h with 2 μM imatinib (IM), 2.5 μM SHIN1, or a combination of both in medium containing 140 μM ^13^C_3_^15^N_1_-serine. (n=4 independent wells from individual experiment). **d**, Mass isotopologue distribution in GSG and GSSG from CML CD34^+^ cells treated for 24 h with 2 μM imatinib, 2.5 μM SHIN1, or a combination of both in medium containing 140 μM ^13^C_3_^15^N_1_-serine (n=4 patient samples). Each isotopologue is shown as a fraction of the sum of all possible isotopologues. Data are presented as the mean ± s.e.m. P-values were calculated using a repeated measure one-way ANOVA with Dunnett’s multiple comparisons test (**d**).

**Extended data Fig.6**: **SHIN1 selectively targets LSCs in combination with standard CML therapy. a,** Gating strategy of flow cytometry analysis to measure engraftment of CML CD34^+^ cells. **b-d**, Frequency of human CD45^+^ (**b**), CD34^+^ (**c**), and CD34^+^CD38^-^ (**d**) of CML cells from parent at experimental endpoint (n=4 mice for untreated group; n=6 mice for imatinib-treated group; n=7 mice for SHIN1-treated group, n=4 mice for combination group). Data are presented as the mean ± s.e.m. P-values were calculated with ordinary one-way ANOVA with Dunnett’s multiple comparisons (**d**).

