## Supplementary data for "Mitochondrial folate metabolism inhibition drives differentiation through mTORC1 mediated purine sensing"

Extended Data Fig. 1:

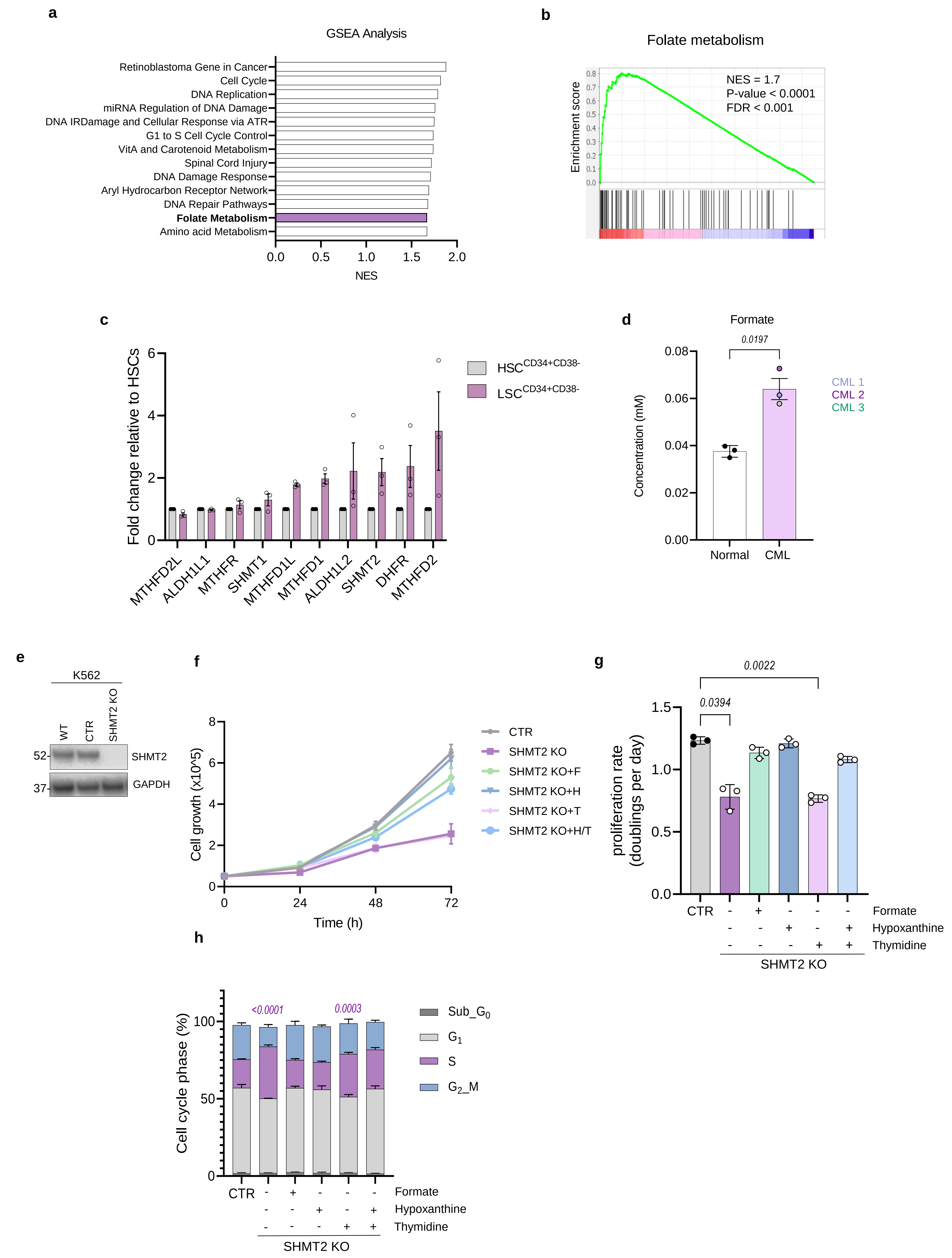

Extended Data Fig. 2:

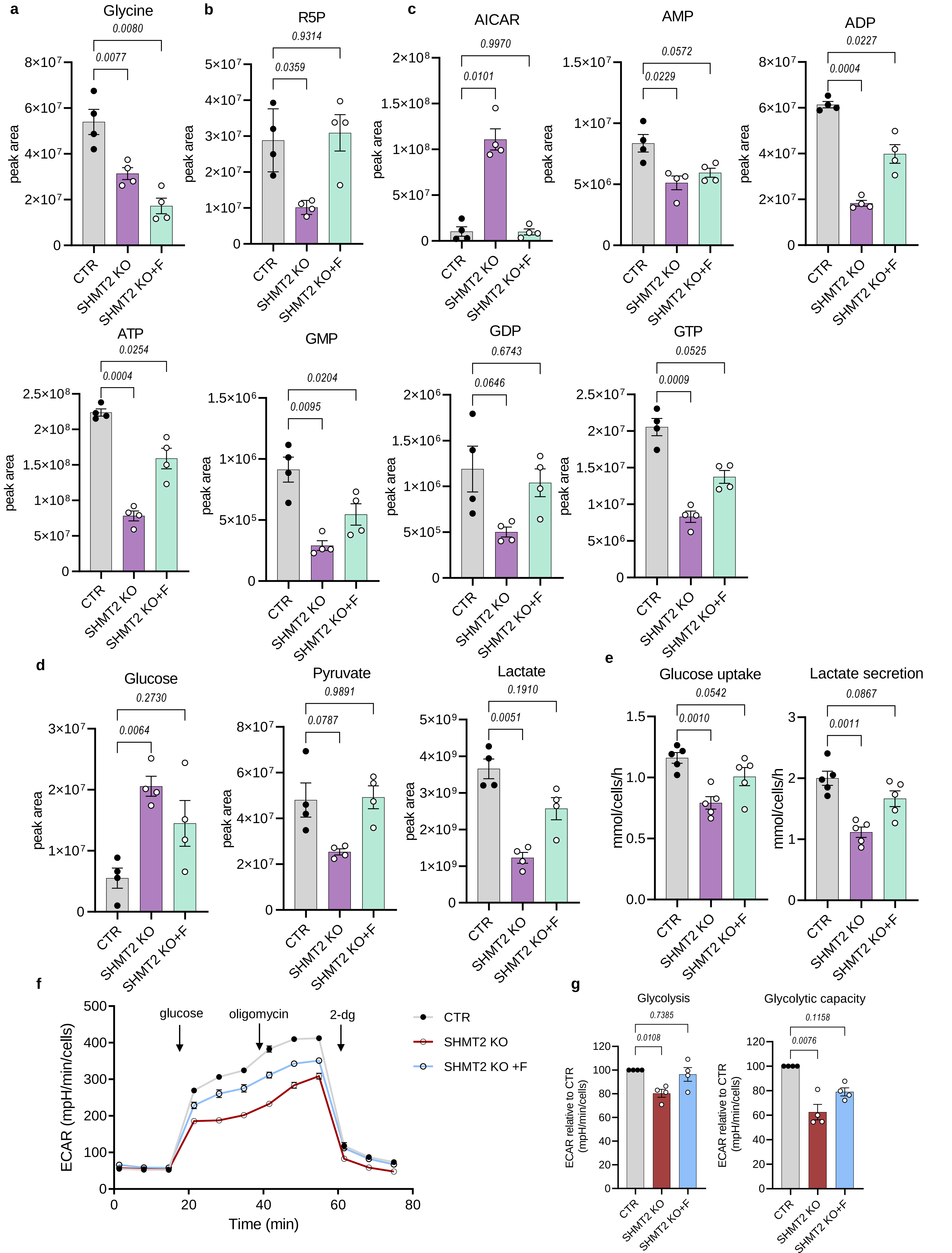

Extended Data Fig. 3:

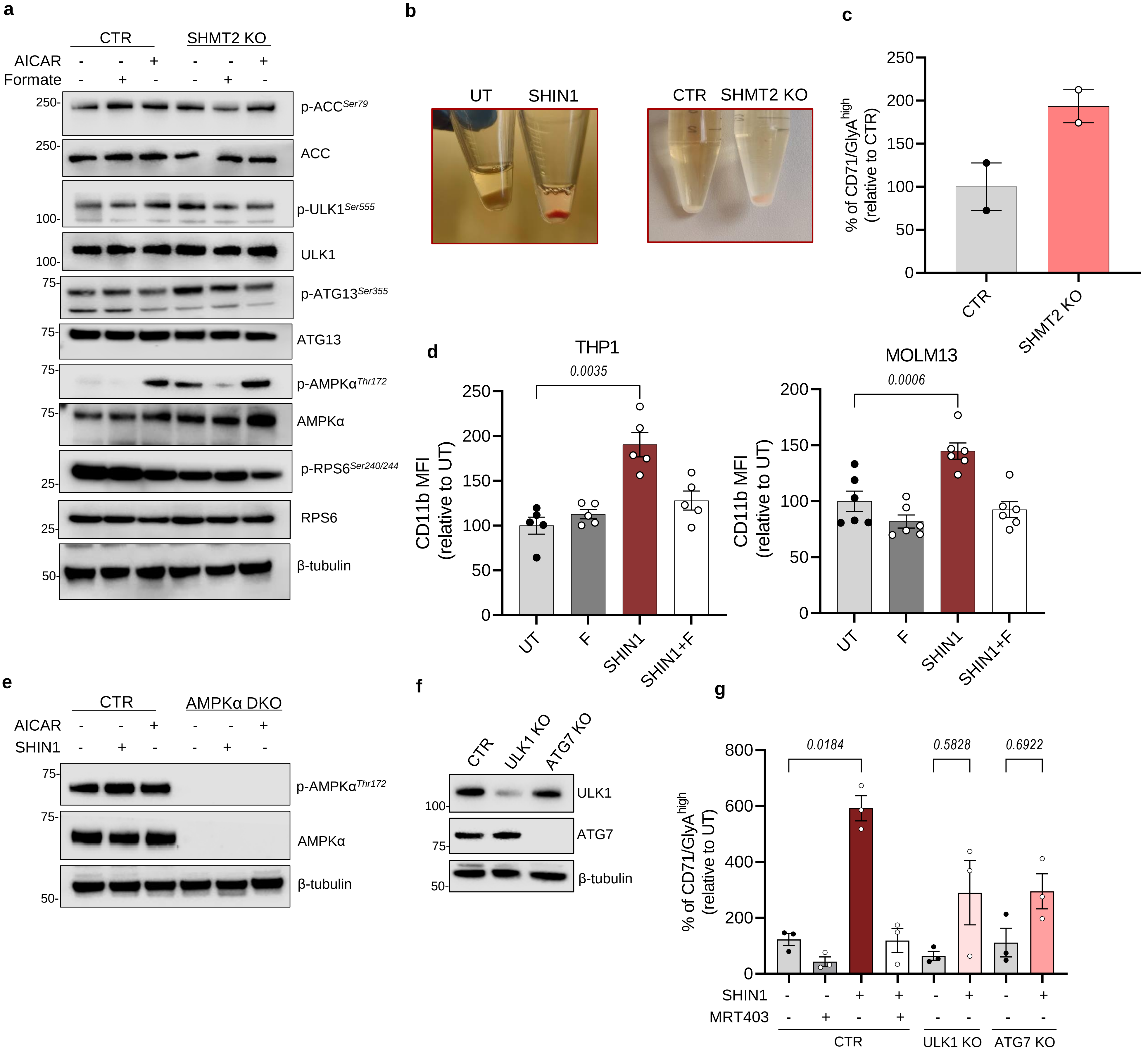

Extended Data Fig. 4:

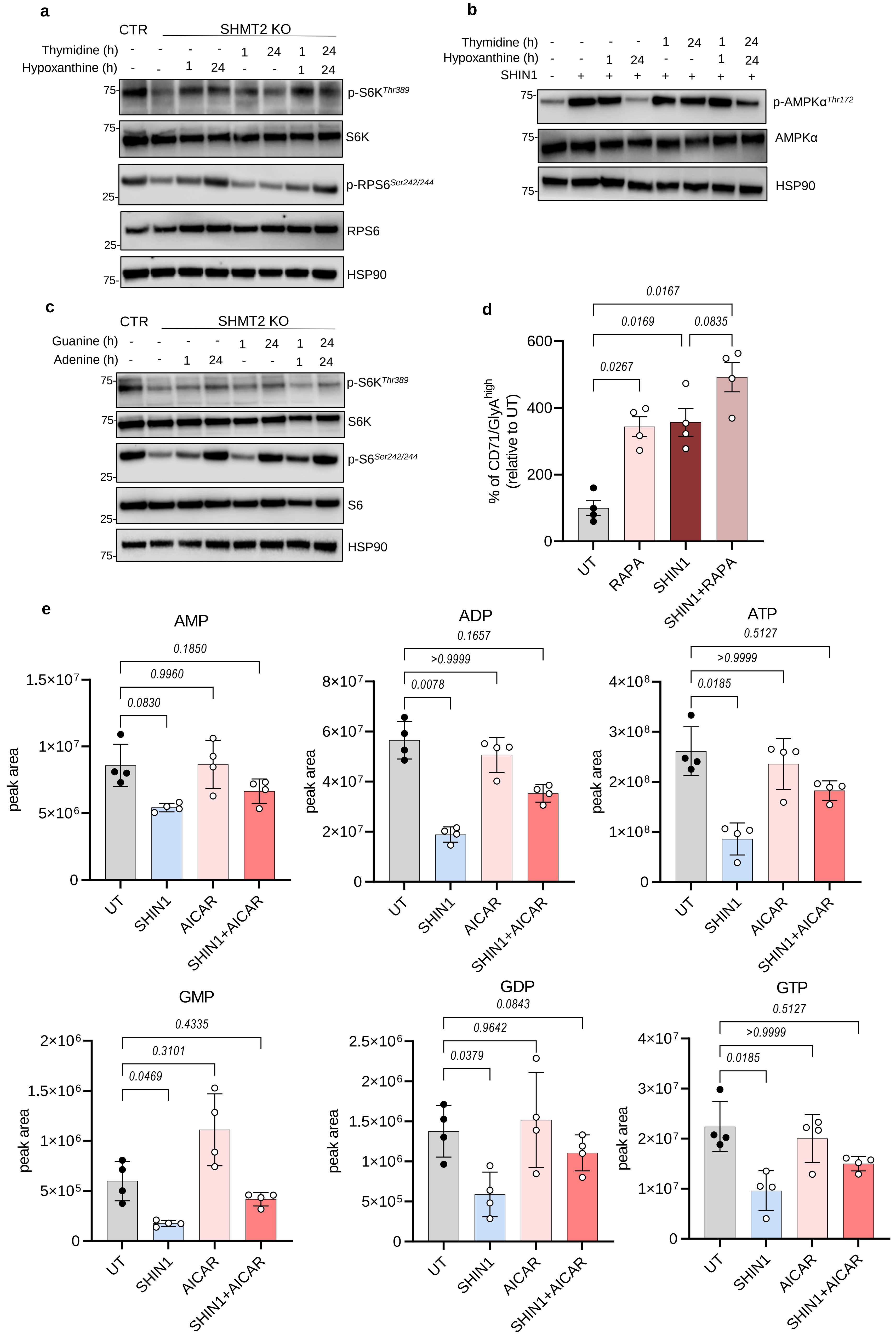

### Extended Data Fig. 5:

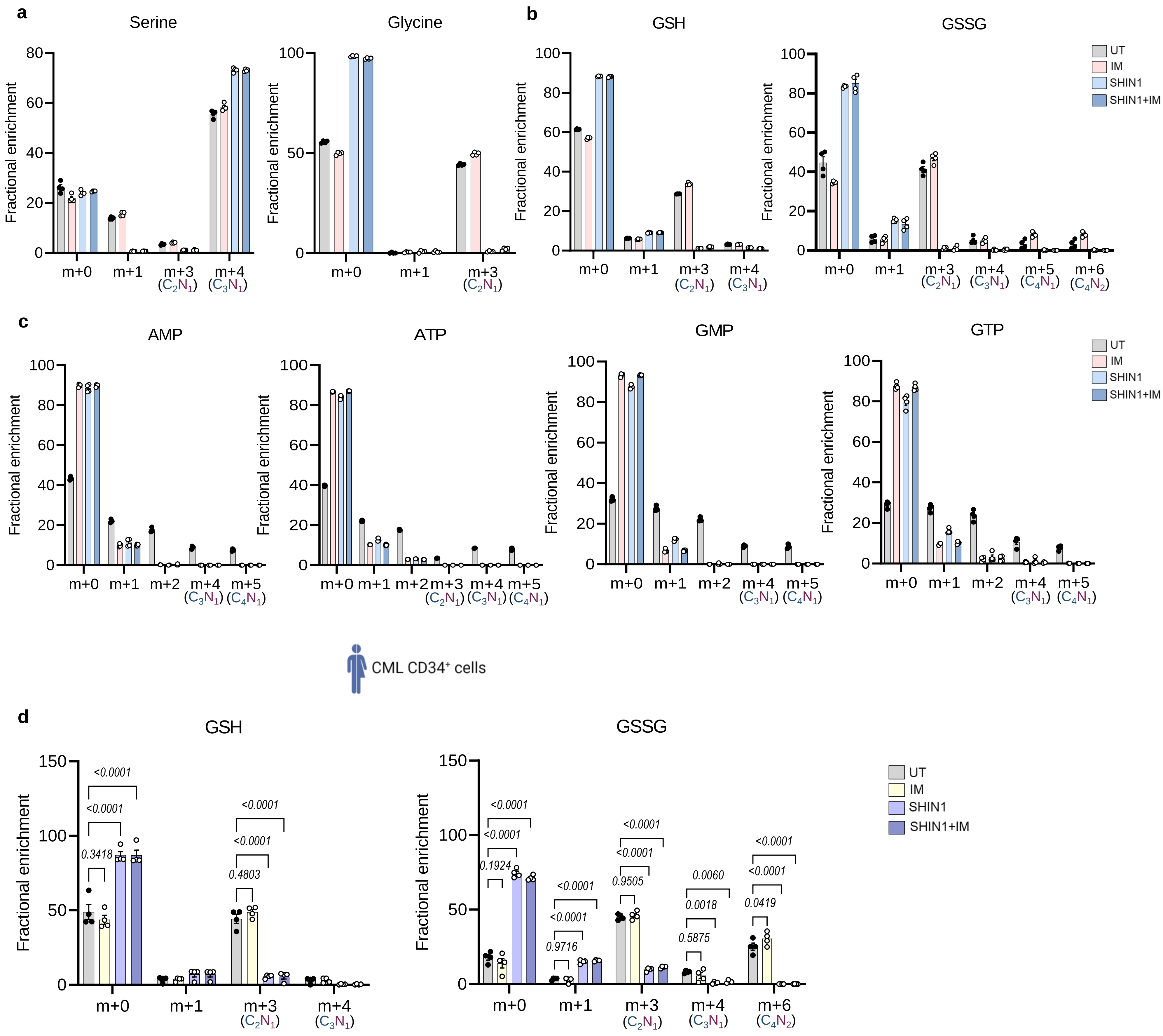

Extended Data Fig. 6:

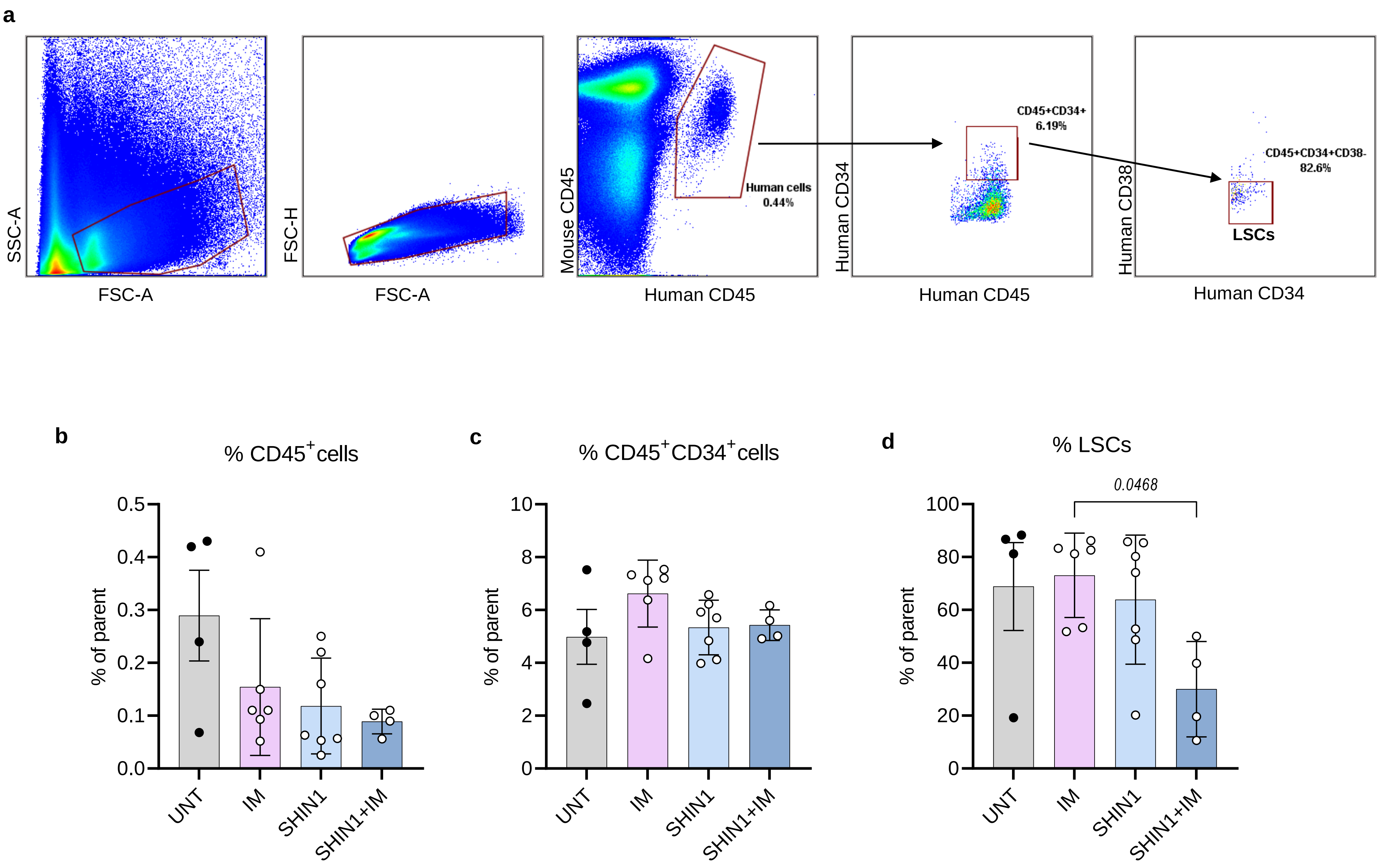
